# Trade-offs between photosynthetic capacity, mesophyll conductance stability and leaf anatomy shape heat and water deficit resilience in *Gossypium*

**DOI:** 10.64898/2026.02.26.708164

**Authors:** Demi Sargent, Warren Conaty, Kelly Chapman, Garima Dubey, Laurel George, Sue Lindsay, Richard Wuhrer, Susanne von Caemmerer, John Evans, Robert Sharwood

**Affiliations:** Hawkesbury Institute for the Environment, Western Sydney University, Richmond, NSW 2753, Australia; CSIRO Agriculture and Food, Narrabri, NSW 2390, Australia; Global Centre for Land-Based Innovation, Hawkesbury Campus, Western Sydney University, Richmond, NSW 2753, Australia; Research School of Biology, Australian National University, Linnaeus Way, Canberra, ACT 2601, Australia; Advanced Materials Characterisation Facility (AMCF), Western Sydney University, Parramatta, NSW 2116, Australia; Macquarie Analytical and Fabrication Facility (MAFF), Macquarie University, Macquarie Park, NSW 2109, Australia

**Keywords:** CO_2_ diffusion, CO_2_-fixation, cotton, drought stress, *Gossypium*, heat stress, mesophyll conductance, photosynthesis

## Abstract

- Mesophyll conductance (*g*_m_) governs CO_2_ diffusion to Rubisco and is a key determinant of photosynthetic performance, yet the mechanisms underlying its sensitivity to heat and water stress remain unresolved.
- We quantified *g*_m_ temperature responses across diverse *Gossypium* species and examined anatomical drivers of *g*_m_ plasticity in cultivated cotton (*G. hirsutum*) and the wild Australian species *G. bickii* under elevated temperature and soil water deficit.
- Species exhibited contrasting *g*_m_ strategies: *G. hirsutum* exhibited high *g*_m_ and carbon assimilation near thermal optima but showed greater sensitivity under combined heat and water deficit, whereas *G. bickii* maintained comparatively stable *g*_m_ and photosynthesis across stress conditions.
- Under water deficit, structural adjustments in *G. hirsutum* (increased leaf porosity, cell wall thickness and mesophyll surface exposure to intercellular airspaces) were insufficient to sustain *g*_m_, suggesting that liquid-phase resistances impose dominant constraints on CO_2_ diffusion under extreme climatic stress.
- These results identify *g*_m_ as a dynamic, multi-component trait and a key physiological vulnerability in cotton, shaped by coordinated anatomical characteristics and potentially cell wall properties and membrane-associated processes, with major implications for mechanistic photosynthesis modelling and improving climate resilience in cotton and other C_3_ species.

## Introduction

Photosynthetic carbon assimilation underpins plant growth, ecosystem productivity, and crop yield. However, it is highly sensitive to environmental stressors, including heat and water limitation. Reductions in photosynthetic rate and efficiency contribute to substantial yield losses during extreme climatic events such as heatwaves and droughts (Cai et al., 2010, Cohen et al., 2021, Fahad et al., 2017, Flexas et al., 2002, Liu et al., 2025). In C_3_ species, photosynthetic limitations arise from stomatal (*L*_s_), mesophyll (*L*_m_) and biochemical (*L*_b_) constraints, where *L*_s_ describes the restriction of CO_2_ entry from the atmosphere into the substomatal cavity or intercellular airspace, and *L*_m_ reflects the resistance to CO_2_ diffusion from the intercellular airspace to the chloroplast stroma (Evans and von Caemmerer, 1996, Flexas et al., 2004, Flexas et al., 2008, Flexas et al., 2012, Grassi and Magnani, 2005, Terashima et al., 2006, Terashima et al., 2011, Tosens et al., 2016, Veromann-Jürgenson et al., 2017).

The prevailing limitation on photosynthesis under environmental stress varies across species (Ennahli and Earl, 2005, Galmés et al., 2007a), light environment (Monneveux et al., 2003, Niinemets and Valladares, 2004), plant age (Varone et al., 2012), and stress history (Flexas et al., 2009, Galle et al., 2011). The dominant limitation also shifts with stress severity. Under mild-moderate water stress, declines in stomatal conductance (*g*_s_) and mesophyll conductance (*g*_m_) typically constrain CO_2_ diffusion and photosynthesis (Cano et al., 2013, Galmés et al., 2007a, Perez-Martin et al., 2014). Under more severe water stress, responses diverge; some studies report pronounced biochemical restriction to assimilation (Chaves et al., 2009, Flexas et al., 2004), while others attribute over 50% of photosynthetic limitation to decreases in *g*_m_ (Perez-Martin et al., 2014).

Photosynthetic responses also depend strongly on stress duration. Rapid responses to water limitation occur within minutes to hours, including stomatal closure followed by decreased *g*_m_ and electron transport (Zou et al., 2022). In contrast, long-term water stress acclimation typically involves changes to leaf anatomy and ultrastructure (Flexas et al., 2006a, Lambers et al., 2008) such as increased leaf thickness and density, altered palisade and spongy tissue ratios and cell rearrangement (Ennajeh et al., 2010, Galmés et al., 2013, Haffani et al., 2017, Han et al., 2019, Tosens et al., 2012a, Zhang et al., 2015). Temperature modulates these responses (Galle et al., 2009); high temperatures frequently exacerbate drought effects (Carmo-Silva et al., 2012, Shah and Paulsen, 2003, Wang et al., 2010), although heat acclimation can reduce susceptibility during combined heat and water stress events (Centritto et al., 2011, Killi et al., 2017). Despite their agronomic, ecological and economic importance, the mechanisms constraining photosynthesis under combined heat and drought stresses that frequently co-occur and may impose antagonistic diffusion and metabolic pressures remain poorly understood (Cai et al., 2015, Cornic, 2000, Lawlor, 1995, Lawlor and Cornic, 2002, Flexas et al., 2009, Flexas and Medrano, 2002, Nadal and Flexas, 2018, Tezara et al., 1999, Zhu et al., 2021).

Mesophyll conductance (*g*_m_) has emerged as a critical bottleneck to photosynthetic capacity, particularly under environmental stress events (Gago et al., 2020). By regulating CO_2_ diffusion from substomatal cavities to chloroplasts, *g*_m_ strongly influences the chloroplastic CO_2_ partial pressure (*C*_c_) and, consequently, Rubisco activation and carboxylation rates. Increasing *g*_m_ can enhance *C*_c_ without increasing *g*_s_, providing a potential strategy for improving both photosynthetic rate and water-use efficiency (Evans and Vellen, 1996, Flexas et al., 2008, Flexas et al., 2012, Flexas et al., 2016, Lauteri et al., 1997, Warren et al., 2007). Substantial genotypic and interspecific variation in *g*_m_ has been documented for C_3_ plants such as rice (*Oryza sativa*, *Oryza glaberrima*), wheat (*Triticum aestivum*), soybean (*Glycine max*), sunflower (*Helianthus annuus*) and cotton (*G. hirsutum*) (Flexas et al., 2008, Gu et al., 2012, Huang et al., 2022, Jahan et al., 2014, Knauer et al., 2019, Ouyang et al., 2017, von Caemmerer and Evans, 2015). Despite its variability, and clear influence on *C*_c_ and modelled photosynthetic parameters such as maximal carboxylation (*V*_cmax_) and electron transport capacity (*J*_max_), *g*_m_ remains underrepresented in crop, ecosystem and carbon-cycle models, where it is often treated implicitly rather than explicitly parameterised (Bernacchi et al., 2002, Flexas et al., 2006a, Flexas et al., 2007, Flexas et al., 2008, Nadal et al., 2021, Sargent et al., 2024, Sharkey et al., 2007, Walker et al., 2013, Warren, 2008b).

The sensitivity of *g*_m_ to environmental factors including light intensity (Galle et al., 2009, Hanba et al., 2002, Laisk et al., 2005, Piel et al., 2002), CO_2_ concentration (Flexas et al., 2007), temperature (Evans and von Caemmerer, 2013, von Caemmerer and Evans, 2015) and water availability (Gu et al., 2012, Jones, 1973) is well established, yet the underlying mechanisms and associated gene networks driving such variation remain unclear (Evans, 2021, Flexas et al., 2008). Short-term temperature response studies reveal highly variable patterns among species. Some exhibit increasing *g*_m_ with temperature until a plateau, others a peaked response with decline at high temperatures, and some show little temperature sensitivity (Bernacchi et al., 2002, Scafaro et al., 2011, von Caemmerer and Evans, 2015, Yamori et al., 2006). Acclimation to growth temperature also differs among species, indicating diverse thermal plasticity of internal CO₂ diffusion (Warren, 2008a). *g*_m_ almost universally declines under water stress (Delfine et al., 2005, Ennahli and Earl, 2005, Flexas et al., 2002, Flexas et al., 2004, Flexas et al., 2006a, Galmés et al., 2007a, Grassi and Magnani, 2005, Monti et al., 2006), although acclimation and recovery after water stress have been observed (Galle et al., 2009, Galle et al., 2011). Additionally, *g*_m_ stress responses vary with stress duration and intensity, and species tolerance (Delfine et al., 2001, Monti et al., 2006). These findings confirm that *g*_m_ is finite, dynamic, and responsive across temporal scales, but the mechanistic basis of this variability, especially under combined heat and water stress, remains unresolved.

Mesophyll conductance reflects resistance along a complex diffusion pathway (Evans, 2021, Salesse-Smith and Xiao, 2025). After entering the leaf through the stomata, gaseous CO_2_ must then diffuse through the intercellular airspace to an exposed mesophyll cell surface, where it then dissolves in the water-filled pores of the cell wall (Evans et al., 2009, Tosens et al., 2012a). Now in the liquid phase, where CO_2_ diffusion is 10,000 times slower in water than in air (Evans et al., 1994, Evans and von Caemmerer, 1996, Terashima et al., 2006), CO_2_ then diffuses through the cell wall, the plasma membrane, cytoplasm, chloroplast envelope and through the chloroplast stroma to reach the active site of Rubisco (Evans et al., 2009, Evans, 2021, Gago et al., 2020, Salesse-Smith and Xiao, 2025, Terashima et al., 2006, Terashima et al., 2011, Tomás et al., 2013, Tosens et al., 2012b, Veromann-Jürgenson et al., 2017).

The main anatomical determinants of *g*_m_ include mesophyll cell wall thickness (*T*_CW_), intercellular airspace fraction (*F*_ias_), chloroplast surface area exposed to intercellular airspace (*S*_c_), and mesophyll surface area exposed to intercellular airspace (*S*_mes_). Among these, *T*_CW_ and *S*_c_ have been identified as key determinants of *g*_m_ responses to environmental factors (Evans et al., 1994, Evans et al., 2009, Nadal et al., 2021, Terashima et al., 2011, Tomás et al., 2013). Mesophyll conductance is typically correlated positively with *S*_c_ and negatively with *T*_CW_ (Evans et al., 2009, Flexas et al., 2021, Terashima et al., 2011, Tosens et al., 2012b, Xiong et al., 2017), which indicates that both the surface area of mesophyll cell walls and their thickness influence *g*_m_.

However, these structural traits do not fully explain the rapid responses of *g*_m_ to short-term temperature and water stress (Flexas et al., 2008), eluding to the influence of aquaporin-mediated CO_2_ transport and carbonic anhydrase-facilitated CO_2_/HCO_3_^-^ interconversion in dynamic regulation of *g*_m_ (Bernacchi et al., 2002, Ermakova et al., 2021, Evans et al., 2009, Evans, 2021, Flexas et al., 2006b, Flexas et al., 2012, Flexas et al., 2021, Flexas et al., 2008, Gago et al., 2020, Gillon and Yakir, 2000b, Gillon and Yakir, 2000a, Groszmann et al., 2017, Hanba et al., 2004, Miyazawa et al., 2008, Momayyezi et al., 2020, Perez-Martin et al., 2014, Price et al., 1994, Terashima et al., 2011, Warren, 2008b). Although the role of aquaporins in regulating *g*_m_ is relatively well supported (Bernacchi et al., 2002, Flexas et al., 2006b, Flexas et al., 2008, Flexas et al., 2012, Galmés et al., 2007b, Han et al., 2016, Hanba et al., 2004, Miyazawa et al., 2008, Perez-Martin et al., 2014), evidence for the influence of carbonic anhydrase is varied (e.g., Han et al. (2016), Han et al. (2019)). More recently, cell wall composition, including pectin content and microfibril alignment, has also emerged as a potential modulator of *g*_m_, particularly under drought (Flexas et al., 2021, Pathare et al., 2024, Roig-Oliver et al., 2022, Roig-Oliver et al., 2025, Salesse-Smith and Xiao, 2025, Sun et al., 2025a, Sun et al., 2025b, Tosens et al., 2012a, Tosens et al., 2012b).

Together, previous research has revealed that *g*_m_ is finite, dynamic and multidimensional, integrating anatomical, biochemical and biophysical controls. Yet the relative contribution of these components underpinning *g*_m_ responses to combined heat and water stress remains unresolved. This knowledge gap constrains both mechanistic understanding of photosynthetic limitations and the ability to predict plant responses under future climate scenarios (Bahar et al., 2018, Evans, 2021, Knauer et al., 2019, Knauer et al., 2020, Nadal et al., 2021, Rogers et al., 2017, Sargent et al., 2024).

The *Gossypium* (cotton) genus provides a unique system to address this knowledge gap. It comprises >50 species adapted to contrasting climates from arid central Australia to humid tropical regions (Fryxell, 1971, Fryxell, 1992, Wendel and Grover, 2015). In several species, domestication has substantially altered leaf anatomy, including enlarged mesophyll cells and reduced cell wall thickness, contributing to higher photosynthetic rates in cultivated cotton (Lei et al., 2022a, Lei et al., 2022b, Lei et al., 2024, Sun et al., 2025b). While *g*_m_ responses to drought (Han et al., 2016, Han et al., 2019), heat (Huang et al., 2022, Mercado Alvarez et al., 2022), and dynamic water stress (Zou et al., 2022) have been reported across *Gossypium*, how anatomical and physiological drivers of *g*_m_ interact under simultaneous heat and water limitation has not been resolved.

To address this, we screened phylogenetically and climatically diverse *Gossypium* species to quantify variation in photosynthetic and *g*_m_ thermal responses, then examined mechanistic drivers in a wild Australian species (*Gossypium bickii*) and a domesticated cultivar (*Gossypium hirsutum* cv. Sicot 71) exposed to elevated growth temperature and severe, long-term soil water deficit. Using carbon isotope discrimination to estimate *g*_m_ in combination with detailed anatomical analyses, we aimed to disentangle the determinants of *g*_m_ and photosynthetic resilience. We hypothesised that species’ climate-of-origin would influence adaptive leaf traits and confer greater *g*_m_ stability in species native to hotter, drier environments. Our findings reveal key mechanisms underpinning *g*_m_ plasticity under combined heat and water stress, providing a basis for improving predictions of photosynthetic limitation and resilience in a warming, drying world.

## Materials and Methods

### Plant growth

All *Gossypium* seed were acid de-linted using 70% sulphuric acid for 2-4 minutes depending on fuzz density, and all “wild” *Gossypium* seed were heat treated in an 80°C water bath for 2 minutes prior to germination on moistened filter paper and transplantation into soil. Seed germination was staggered to accommodate for different growth rates between species and to enable measurements to be taken at the same stage of phenological development. All plants were grown in a commercial premium potting mix (Martins premium potting mix) in 10L pots. To ensure plant growth was nutrient-replete, slow-release fertiliser (Scotts Osmocote Tomato, Vegetable and Herb Controlled Release Fertiliser) was applied once at sowing at 15 g per pot, and soluble fertilizer (Yates Thrive Flower and Fruit) was applied fortnightly at 4 g L^-1^, with 1L of solution supplied per pot. During the soil water deficit treatment, nutrient supply was maintained by dissolving 2 g of the soluble fertiliser into the reduced irrigation volume for weekly application to water deficit treated plants, ensuring equivalent nutrient supply per plant across watering treatments and avoiding confounding effects of nutrition. The cultivar used in these experiments, ‘Sicot 71’, is considered representative germplasm of a standard conventional commercial Australian cotton cultivar.

### Temperature response experiment

Seed of four diverse diploid *Gossypium* species from diverse geographical origins (*G. arboreum* from Asia, *G. bickii* and *G. sturtianum* from Australia and *G. gossypioides* from Oaxaca, Mexico) and a tetraploid cotton cultivar (*G. hirsutum* cv. Sicot 71) were sown from 12^th^ September 2022 to 24^th^ October 2022. These plants were grown in a temperature-controlled glasshouse under supplemental light (Mars Hydro full spectrum LED grow lights) at the Hawkesbury Institute for the Environment, Western Sydney University, Richmond, NSW. Glasshouse air temperature was controlled to 32°C (day) and 21°C (night), and plants were watered to soil saturation every 1-3 days to ensure adequate hydration.

At “squaring” (first appearance of immature floral buds), the temperature response of mesophyll conductance (*g*_m_) and net leaf photosynthesis (*A*_380_) was measured on the first fully expanded leaf of each plant (*n*=5) across four temperatures (25, 30, 35 and 40°C), as detailed in Sargent et al. (2024). For these temperature response measurements, plants were transferred from the glasshouse into a temperature-controlled growth cabinet (Thermoline Climatron; Thermoline Scientific, Sydney, NSW, Australia) to enable precise control of measurement temperature. Air temperature within the cabinet was set to each treatment temperature, and leaf temperature was matched to the target setpoints using the LI-6400XT block temperature control, ensuring close agreement between air and leaf temperatures during gas exchange measurements. Measurements were made with the sample CO_2_ maintained at 380 µmol CO_2_ mol^-1^ air, according to von Caemmerer and Evans (2015) and Evans and von Caemmerer (2013).

### Soil water deficit (SWD) and elevated temperature experiment

For the elevated temperature and soil water deficit experiment, seed of *G. bickii* native to semi-arid regions of Australia and a cotton cultivar (*G. hirsutum* cv. Sicot 71) were sown on 27^th^ July 2022 and 11^th^ August 2022, respectively. These plants were grown in a temperature-controlled glasshouse at the Hawkesbury Institute for the Environment, Western Sydney University, Richmond, NSW, under supplemental light (Mars Hydro full spectrum LED grow lights) at either control (32°C (day)/21°C (night)) or elevated (at 38°C (day)/27°C (night)) temperatures. At 9 to 11 weeks after germination for *G. hirsutum* and *G. bickii*, respectively, the soil water deficit treatment was applied for approximately six weeks.

Soil water content was measured gravimetrically using pot balances. “Well-watered” (WW) pots were watered to full saturation of the soil every 1-3 days. Water was withheld from “water deficit” (WD) pots from 17^th^ October 2022, and the soil water content of individual pots was maintained at 60-75% soil water deficit (SWD) for approximately six weeks by applying gravimetrically measured amounts of water to each pot every 1-3 days (Fig. 1). Plants were monitored for wilting, and leaf water potential (LWP; Fig. 2A) was measured using a pressure chamber periodically to ensure adequate SWD levels.

**Figure 1.**
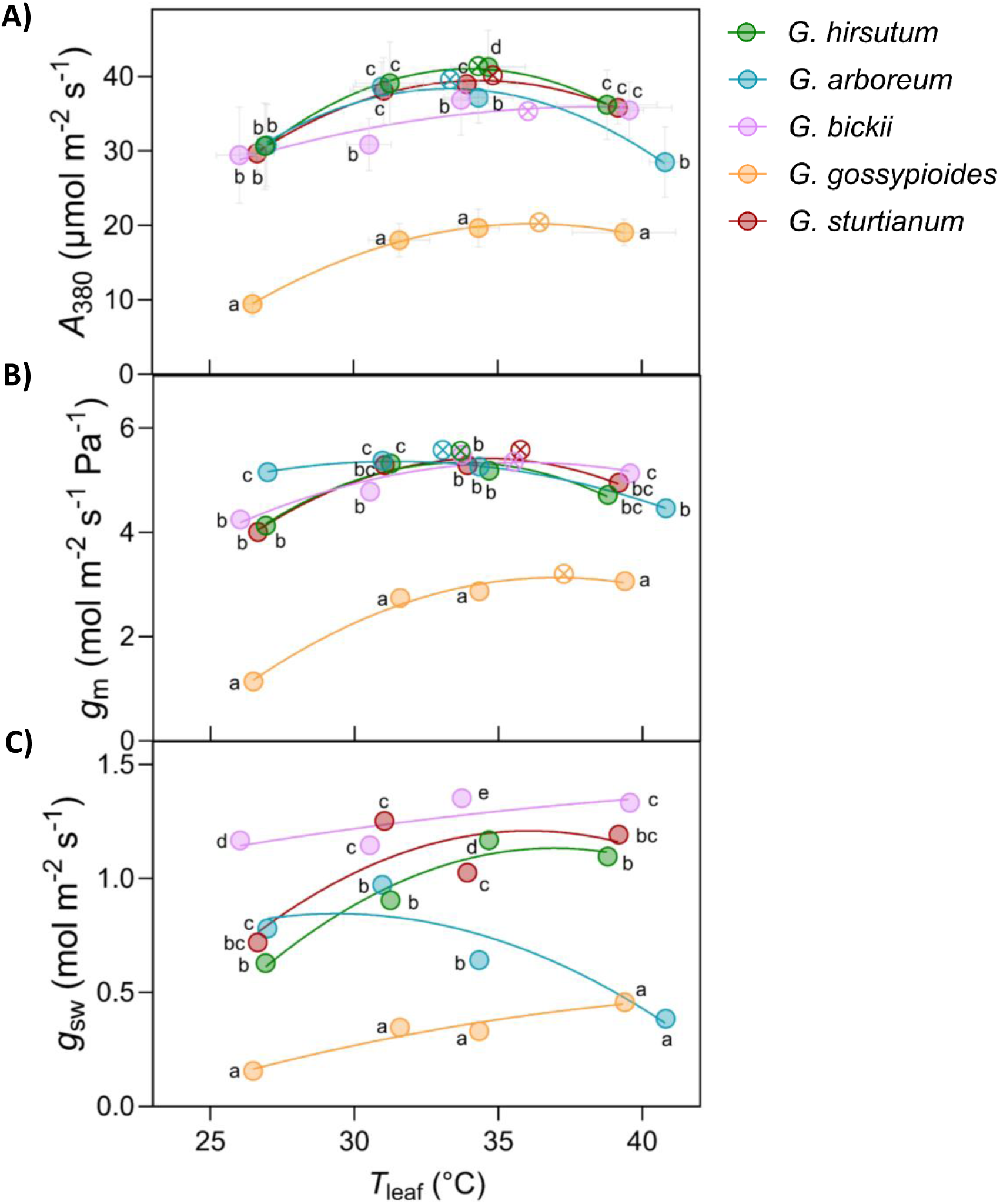
Temperature responses of A) net leaf photosynthesis measured at 380 µmol CO2 mol^-1^ air (*A*380), B) mesophyll conductance to CO2 (*g*m), and C) stomatal conductance to water (*g*sw) in *G. hirsutum* cv. Sicot 71 (green), *G. arboreum* (blue), *G. bickii* (pink), *G. gossypioides* (orange) and *G. sturtianum* (red). Values are means of five replicate plants. Mean thermal optima for *g*m and *A*380 for each species are indicated by quartered circle symbols. Solid lines show second-order polynomial fits to the mean data. Error bars represent ±SD. Different letters indicate significant differences among species within a given temperature (Fisher’s protected LSD, *p*<0.05).

**Figure 2.**
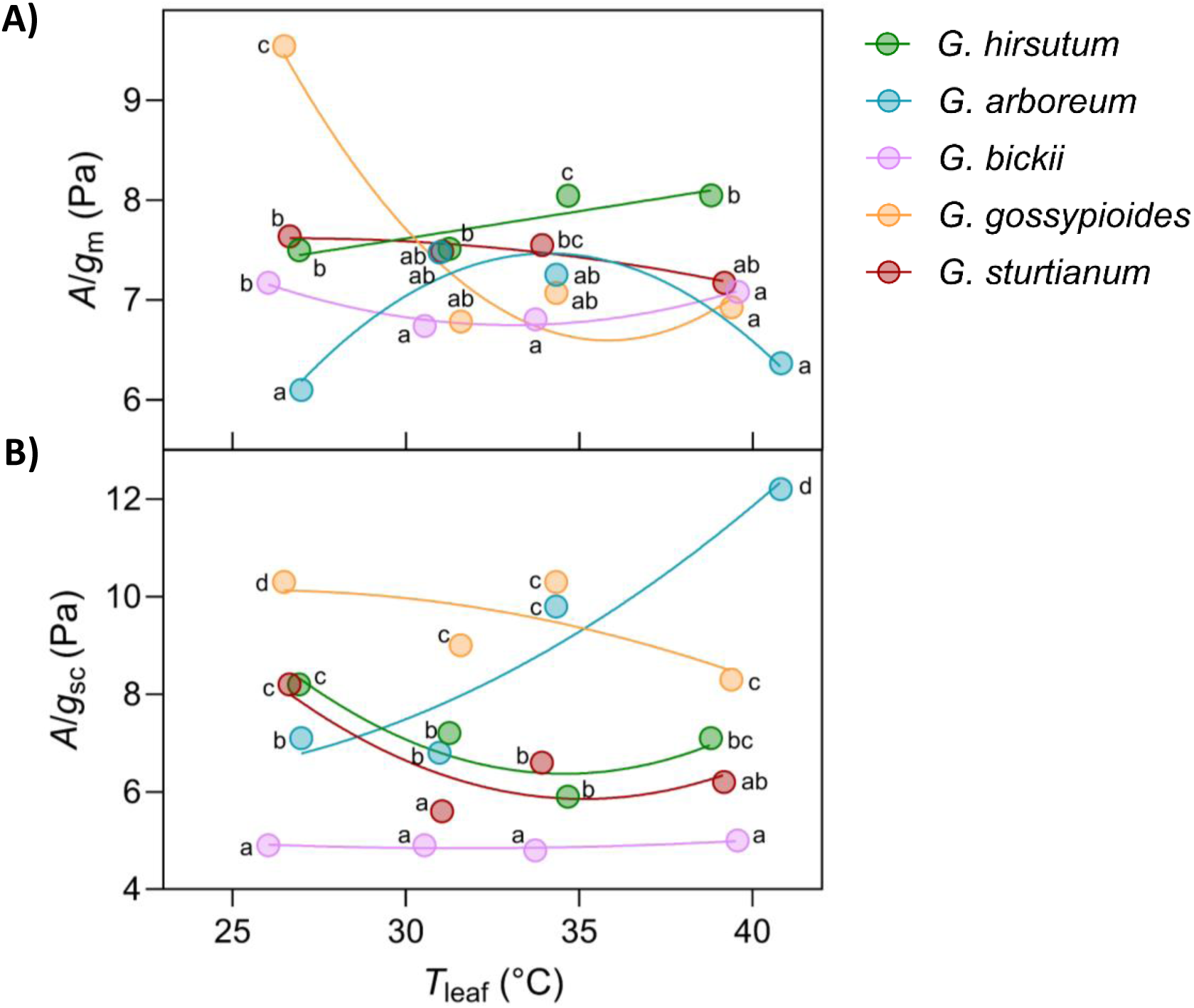
Temperature responses of A) CO2 drawdown from the substomatal cavities to the sites of carboxylation within the chloroplasts (*A*/*g*m), and B) CO2 drawdown from the ambient air to the substomatal cavities (*A*/*g*sc) in *G. hirsutum* cv. Sicot 71 (green), *G. arboreum* (blue), *G. bickii* (pink), *G. gossypioides* (orange) and *G. sturtianum* (red). Values represent means of five replicate plants. Solid lines show second-order polynomial fits to the mean data. Different letters indicate significant differences among species within a given temperature (Fisher’s protected LSD, *p*<0.05)

After six weeks of water deficit treatment, *g*_m_ was measured by carbon isotope discrimination using two LI-6400XT Portable Photosynthesis Systems (LICOR Biosciences Inc., Lincoln, Nebraska, USA) with infra-red gas analyser (IRGA) sensor heads which were coupled to two tunable diode lasers (TDL), as detailed in Sargent et al. (2024). To ensure precise control of measurement temperature, plants were transferred from the glasshouse into a temperature-controlled growth cabinet (Thermoline Climatron; Thermoline Scientific, Sydney, NSW, Australia), where cabinet air temperature was set to match the respective growth temperature treatments during gas exchange and isotope discrimination measurements.

After completion of *g*_m_ measurements, leaf tissue was collected for microscopic analysis from the same region of the leaf enclosed in the gas exchange chamber.

### Measurement of *g*_m_ by carbon isotope discrimination

Carbon isotope discrimination was used to calculate *g*_m_ following Evans and von Caemmerer (2013), with ternary corrections applied according to Farquhar and Cernusak (2012). Two LI-6400XT Portable Photosynthesis Systems (LICOR Biosciences Inc., Lincoln, Nebraska, USA) with infra-red gas analyser (IRGA) sensor heads were coupled to a tunable diode laser (TDL; TGA100, Campbell Scientific, Inc., Logan, UT, USA), which sequentially sampled reference and sample air streams at 4-minute intervals as described by Tazoe et al. (2011) and von Caemmerer and Evans (2015).

For all measurements, plants were placed inside a temperature-controlled growth cabinet (Thermoline Climatron; Thermoline Scientific, Sydney, NSW, Australia), with cabinet air temperature set to match treatment temperatures. Leaf chamber block temperature was manually controlled to maintain leaf temperature (*T*_leaf_, °C) at the target treatment temperature throughout measurements. The LI-6400XT inlet air was supplied with 2% O_2_ to suppress photorespiration and minimise associated fractionation effects on the discrimination signal, enabling more robust estimation of *g*_m_ (Evans and von Caemmerer, 2013; see Appendix 1 of Sargent et al., 2024). Leaf chamber conditions were maintained at 50-70% relative humidity, 200 µmol s^-1^ IRGA flow rate, 380 µmol mol^-1^ air sample CO_2_ concentration, and PAR 2000 µmol m^-2^ s^-1^. Leaf vapour pressure deficit (VPD_L_) was maintained between 1 and 4 kPa. Plants and instruments were allowed to stabilise for ≥30 minutes prior to measurement, and instantaneous gas exchange and carbon isotope discrimination were logged at 4-minute intervals.

### Microscopic analysis of leaf anatomical characteristics

In-tact whole leaves were covered in foil and left overnight to minimise starch accumulation. Leaf tissue was collected between 08:00AM and 10:00AM. From each leaf, three leaf lamina samples were excised as 5mm × 10mm strips using a razor blade and immediately submerged in fixative solution (2.5% glutaraldehyde in 0.1M sodium phosphate buffer, pH 7.2) with 50µL/mL 10% Tween 20. Samples were vacuum infiltrated and fixed overnight at 4°C. Following fixation, the samples were dehydrated through a graded ethanol series, infiltrated with LR White resin and polymerised overnight at 60°C.

For light microscopy, semithin transverse sections (0.5-1.0µm thick) were cut with glass knives using an RMC PT-X PowerTome ultramicrotome and stained with hot aqueous 0.05% Toluidine blue (pH 5.0). Sections were imaged under 10× and 40× objectives using an [insert microscope model and company]. For STEM analysis, ultrathin transverse sections (70-90nm thick) were cut from the same region as semithin light microscopy sections using a diamond knife on the RMC PT-X PowerTome ultramicrotome. Sections were mounted onto carbon film-coated copper EM grids, double stained with Uranyl Acetate Replacement Stain for 1hr and Lead Citrate 3% for 1min and imaged on a Zeiss Merlin field emission gun scanning electron microscope (FEGSEM) operated at 30 kV and using a STEM detector.

Quantitative analysis of leaf anatomical parameters was conducted using ImageJ software (v1.53e; NIH, USA) following the methods described by Evans et al. (1994). Leaf thickness, mesophyll thickness, intercellular airspace fraction (*F*_ias_), and mesophyll surface area exposed to the intercellular airspace (*S*_mes_) were measured from light micrographs. *F*_ias_ was calculated as the proportion of mesophyll area occupied by airspace relative to total mesophyll area. *F*_ias_ and *S*_mes_ were quantified using the “freehand line” tool to trace individual mesophyll cells and intercellular airspace boundaries, and calculated using the “measure” function in ImageJ.

Measurements of leaf thickness, mesophyll thickness and *S*_mes_ were taken from three technical replicates (different sections) across three biological replicates (leaves from different plants). Cell wall thickness (*T*_CW_) was quantified from STEM micrographs on at least nine technical replicates (individual cells) across three biological replicates. Both spongy and palisade were included to account for potential tissue-specific variation.

### Statistical analyses

Analysis of variance (ANOVA) was conducted on each dataset using Genstat v23.0 (VSN International, Hemel Hempstead, UK). Statistical differences were assessed at a 95% confidence level (*p*≤0.05). Species, temperature, water and their interactions were treated as fixed effects. Where ANOVA indicated significant effects, pairwise comparisons were performed using Fisher’s protected least significant difference (LSD) test. All data were checked for normality, linearity and homogeneity prior to analysis.

Regression analyses and ANCOVA/multiple linear regression were performed to test for treatment or group effects, including the influence of continuous covariates such as leaf or mesophyll anatomical traits on physiological responses.

## Results

### Temperature responses of mesophyll conductance between *Gossypium* species

Interspecies variation was evident in the temperature responses of *A*_380_, *g*_m_, *g*_sw_, *A*/*g*_m_ and *A*/*g*_sc_ (Fig. 1 and Fig. 2). *A*_380_ increased with temperature to thermal optima between 33 and 36°C, followed by a decline at high temperatures (Fig. 1A). The temperature at which *A*_380_ peaked differed significantly among species (*p* <0.001), with the highest optima observed in *G. gossypioides* (36.5°C), *G. bickii* (36.1°C) and *G. sturtianum* (34.8°C), while *G. hirsutum* (34.3°C) and *G. arboreum* (33.3°C) exhibited the lowest thermal optima. The peak *A*_380_ value (*p* <0.001) was greatest in *G. hirsutum* (41.4 µmol m^-2^ s^-1^), *G. sturtianum* (40.2 µmol m^-2^ s^-1^) and *G. arboreum* (39.6 µmol m^-2^ s^-1^), followed by *G. bickii* (35.3 µmol m^-2^ s^-1^), with *G. gossypioides* exhibiting the lowest peak (20.4 µmol m^-2^ s^-1^). At supra-optimal temperatures, *G. hirsutum* and *G. arboreum* showed the most pronounced declines in *A*_380_, decreasing by 12.3% and 26.2%, respectively. In contrast, *G. bickii* and *G. gossypioides* exhibited the most thermally stable response, with only modest declines in *A*_380_ of 3.8 and 2.9%, respectively.

Similarly, in all species except one, *g*_m_ increased with temperature, reaching thermal optima between 33 – 36°C, followed by a distinct decline at 40°C (Fig. 1B). The decline in *g*_m_ from 35°C to 40°C was more pronounced for *G. hirsutum* (9 – 11%) compared to the Australian species, *G. bickii* (6%) and *G. sturtianum* (7%). In contrast, *G. arboreum* exhibited a shallow thermal response, with *g*_m_ maintained between 5.2 and 5.4 mol m^-2^ s^-1^ Pa^-1^ from 25 to 35°C, before declining by 13 – 17% to 4.5 mol m^-2^ s^-1^ Pa^-1^ at 40°C. Notably, *g*_m_ for *G. gossypioides* increased strongly with temperature to 3.2 mol m^-2^ s^-1^ Pa^-1^ at a thermal optimum of 37.3°C. However, this species consistently exhibited the lowest *g*_m_ values across all temperatures (*p*<0.001).

Species differed in the temperature response of stomatal conductance to water (*g*_sw_; Fig. 1C), which showed greater variability across temperatures than *A*_380_ and *g*_m_. In *G. bickii* and *G. gossypioides*, *g*_sw_ increased gradually with temperature. *G. hirsutum* and *G. sturtianum* exhibited steeper increases in *g*_sw_ with temperature, followed by a plateau between 35 and 40°C. Uniquely, *G. arboreum* showed a decline in *g*_sw_ with increasing temperature, decreasing from 0.8-1 mol m^-2^ s^-1^ at 25-30°C to 0.4 mol m^-2^ s^-1^ at 40°C. At 25°C, *g*_sw_ was lowest for *G. gossypioides* (0.2 mol m^-2^ s^-1^), intermediate for *G. hirsutum* (0.6 mol m^-2^ s^-1^), *G. sturtianum* (0.7 mol m^-2^ s^-1^) and *G. arboreum* (0.8 mol m^-2^ s^-1^), and highest for *G. bickii* (1.2 mol m^-2^ s^-1^; *p*<0.001). At 40°C, *G. bickii* maintained the highest *g*_sw_ (1.3 mol m^-2^ s^-1^), comparable to *G. sturtianum* (1.2 mol m^-2^ s^-1^) but greater than *G. hirsutum* (1.1 mol m^-2^ s^-1^), while *G. gossypioides* (0.5 mol m^-2^ s^-1^) and *G. arboreum* (0.4 mol m^-2^ s^-1^) exhibited the lowest values (*p*<0.001).

*A*/*g*_m_ represents the CO_2_ drawdown from the substomatal cavities to the sites of carboxylation within the chloroplasts (*C*_c_). Across species, *A*/*g*_m_ exhibited little temperature dependence (Fig. 2A), with the exception of *G. arboreum* which followed a quadratic response, increasing to an apparent optimum before declining at supraoptimal temperatures. At low temperature, *G. gossypioides* exhibited the largest *C*_i_-*C*_c_ drawdown, reflecting its characteristically low *g*_m_ (Fig. 1B), with *A*/*g*_m_ values 25 – 56% higher than those of other species (*p* <0.001).

*A*/*g*_sc_, the CO_2_ drawdown from the ambient air (*C*_a_) to the substomatal cavities (*C*_i_), quantifies the diffusional limitation imposed by stomatal conductance (*C*_a_ – *C*_i_ = *A*/*g*_sc_). There was little temperature dependence for *A*/*g*_sc_ except for *G. arboreum* which showed a strong positive temperature response (Fig. 2B). *A*/*g*_sc_ was broadly similar among species, except at 40°C where it was highest for *G. arboreum* (12.2 Pa; *p* <0.001) followed by *G. gossypioides* (8.3 Pa) and *G. hirsutum* (7.1 Pa), and lowest for *G. bickii* (5.0 Pa) and *G. sturtianum* (6.3 Pa). Across temperatures, *G. bickii* consistently displayed the lowest *A*/*g*_sc_ (4.8 – 5.0 Pa), coinciding with higher *g*_sw_ and *C*_i_ relative to most other species (Table S2).

### Responses of mesophyll conductance and photosynthesis to elevated temperature and soil water deficit differ between *G. hirsutum* and *G. bickii*

Air temperature and soil water content significantly influenced *g*_m_, *A*_380_ and *A*/*g*_m_ (Fig. 3), with differing responses observed between *G. hirsutum* and *G. bickii*. Compared to control conditions (CT_WW: well-watered at 32°C), *g*_m_ decreased in both species under soil water deficit at control temperature (CT_WD), with a more pronounced reduction observed in *G. hirsutum* (−31%) than *G. bickii* (−19%; Fig. 3A). In contrast, under elevated temperature when well-watered (ET_WW), *g*_m_ increased by a similar magnitude in both species (40 – 41%). However, *g*_m_ remained unchanged under combined stresses (ET_WD).

**Figure 3.**
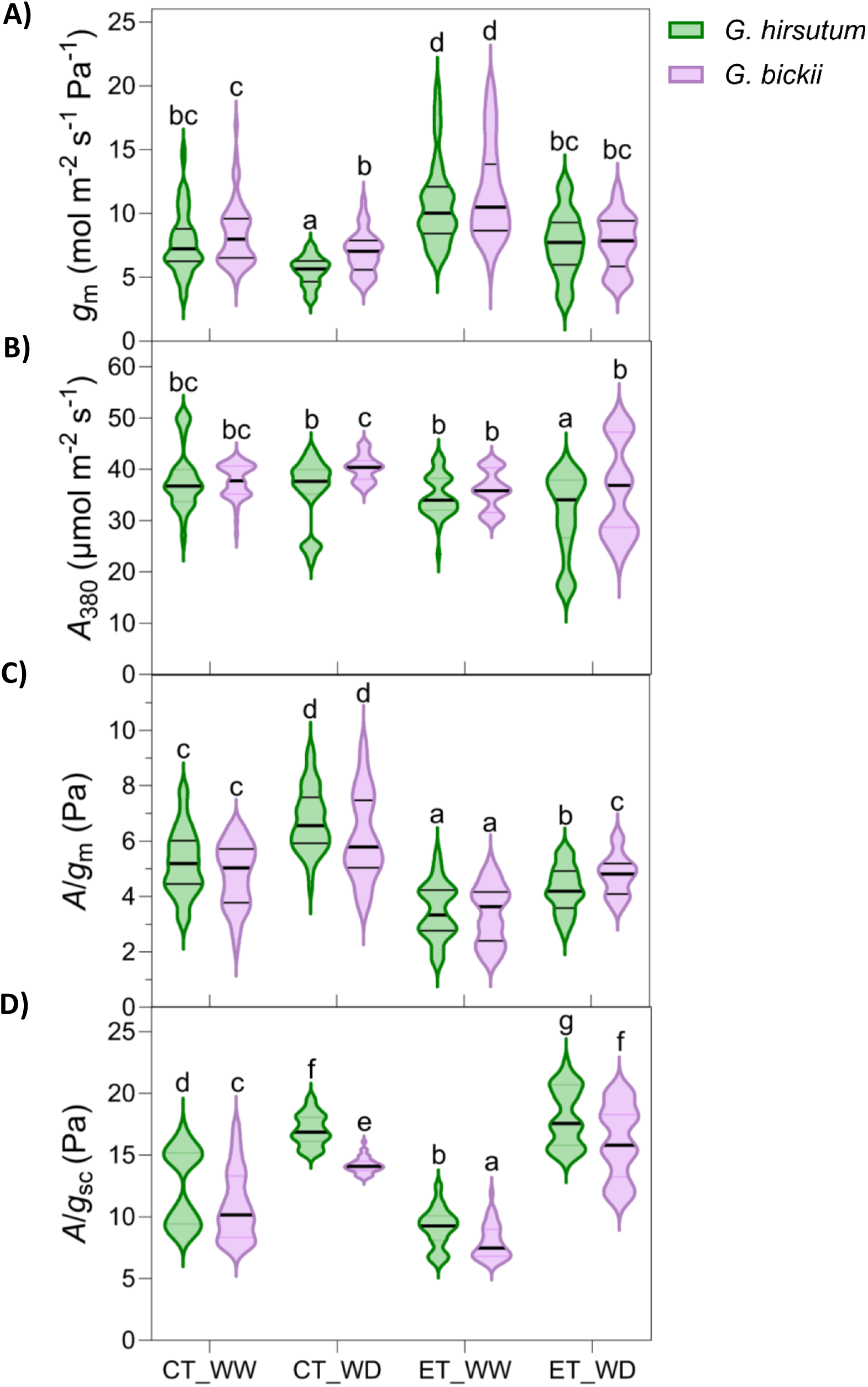
A) Mesophyll conductance (*g*m; *p* <0.001), B) net leaf photosynthesis measured at 380 µmol CO2 mol^-1^ air (*A*380), C) CO2 drawdown from the substomatal cavities to the sites of carboxylation within the chloroplasts (*A*/*g*m), and D) CO2 drawdown from the ambient air to the substomatal cavities (*A*/*g*sc) for *G. hirsutum* (green) and native Australian species *G. bickii* (pink) under different combinations of control (CT) and elevated (ET) temperatures and well-watered (WW) and water-deficit (WD) treatments. Violin plots show the distribution of 60 technical replicates across 6 biological replicates.

*A*_380_ remained largely unchanged across all treatments, except in *G. hirsutum* under combined elevated temperature and soil water deficit (ET_WD), where it declined by 20% (Fig. 3B). Relative to control conditions, *A*/*g*_m_ (Fig. 3C) increased by 29% under soil water deficit (CT_WD) across both species, likely due to the reduction in *g*_m_ (Table S1). Conversely, *A*/*g*_m_ decreased by 31 – 32% under elevated temperature (ET_WW) across both species, consistent with the observed increase in *g*_m_. Both species responded similarly across treatments, except under combined stresses (ET_WD), where *G. hirsutum* exhibited a 17% decline in *A*/*g*m relative to control, while *G. bickii* maintained a stable *A*/*g*m. *A*/*g*sc was influenced by species, water treatment, and temperature treatment, and the interactions between species and water, as well as between water and temperature (*p* <0.001; Fig. 3D). Across all treatments, *G. hirsutum* consistently exhibited higher *A*/*g*sc than *G. bickii*, unique from the *A*/*g*m responses. Within each species, *A*/*g*sc was lowest under elevated temperature with well-watered conditions (ET_WW; 5.7 Pa for *G. hirsutum*, 5.0 Pa for *G. bickii*), intermediate under control well-watered conditions (CT_WW; 7.8 Pa, 6.8 Pa) and water deficit conditions (CT_WD; 10.6 Pa, 8.9 Pa), and highest under combined elevated temperature and water deficit (ET_WD; 11.3 Pa, 10.1 Pa).

### Differing gas exchange capabilities afford potential benefits to wild Australian *G. bickii*

Interspecies differences in gas exchange responses to elevated temperature and soil water deficit were evident between *G. hirsutum* and *G. bickii*. Under well-watered conditions at 32°C, both species exhibited moderately high stomatal conductance (*g*sw; 0.53 – 0.59 mol m^-2^ s^-1^; Fig. 4A). Across all treatments except control conditions, *G. bickii* consistently maintained higher *g*sw than *G. hirsutum* (*p* <0.001). Soil water deficit reduced *g*sw in both species, with *G. hirsutum* experiencing a more pronounced decline: reductions of 38% and 48% at 32°C and 38°C, respectively, compared with 22% and 36% for *G. bickii*. Elevated temperature under well-watered conditions stimulated an increase in *g*sw, more so in *G. bickii* (27%) than in *G. hirsutum* (23%).

**Figure 4.**
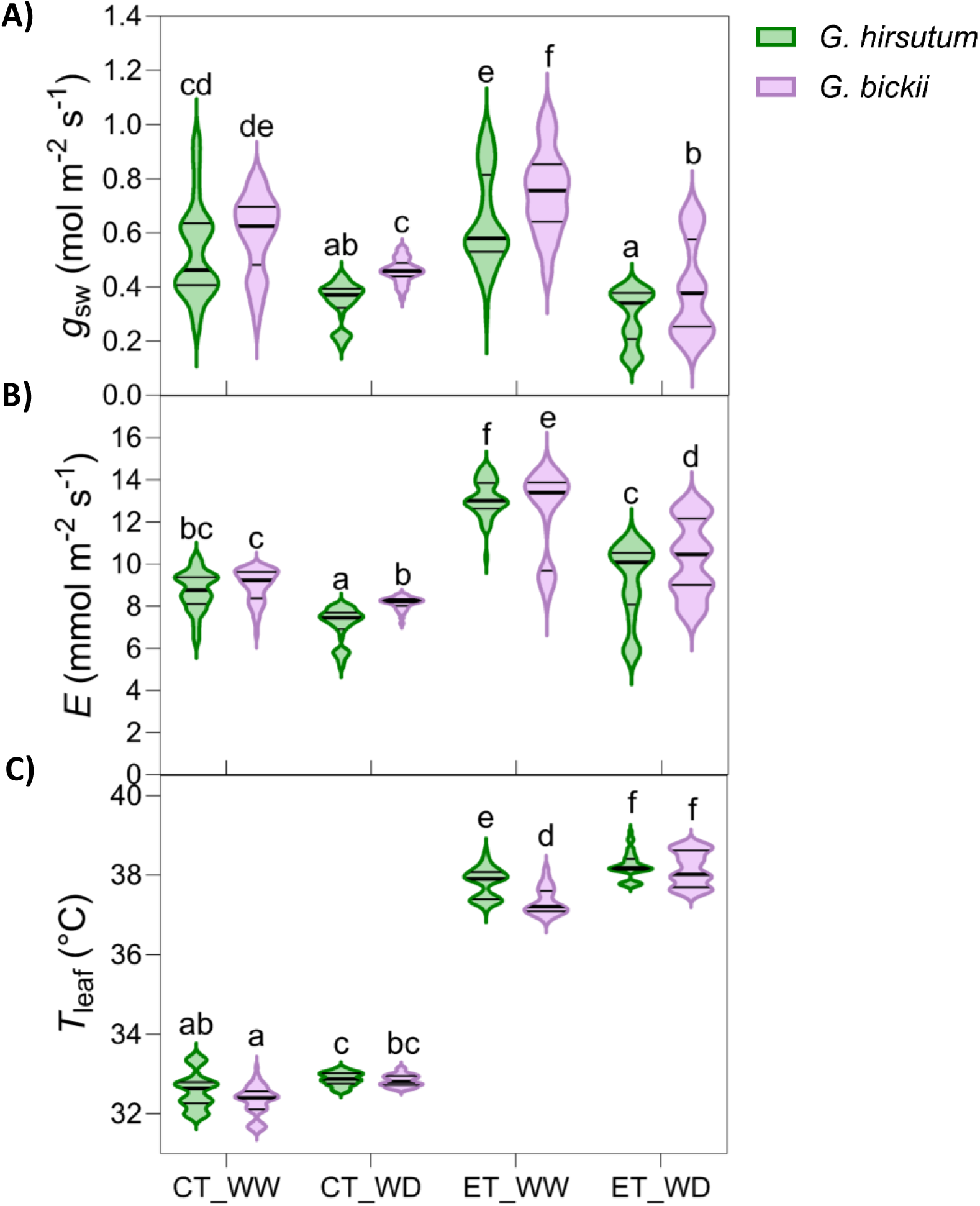
A) Stomatal conductance to water (*g*sw), B) the rate of transpiration of water vapour (*E*), and C) leaf temperature (*T*leaf) for *G. hirsutum* cv. Sicot 71 (green) and *G. bickii* (pink) under different combinations of control (CT) and elevated (ET) temperatures and well-watered (WW) and water-deficit (WD) treatments. Violin plots show the distribution of 60 technical replicates across 6 biological replicates.

Transpiration rate (*E*) followed similar trends to *g*sw, with higher values in *G. bickii* across treatments (Fig. 4B). Elevated temperature under well-watered conditions enhanced *E* by 40 – 52%, while soil water deficit at 32°C reduced *E* by 8 – 20% in both species. However, under soil water deficit at 38°C, *E* was maintained in *G. hirsutum* and increased by 13% in *G. bickii*, diverging from the trend observed in *g*sw.

Leaf temperature (*T*leaf) was consistently higher under water deficit across species, with *T*leaf in CT_WD exceeding CT_WW and ET_WD exceeding ET_WW for both species (Fig. 4C). Species differences in *T*leaf were generally negligible, except under well-watered conditions at 38°C, where *G. bickii* maintained a lower *T*leaf than *G. hirsutum* (−0.42°C, *p* <0.001). This difference coincided with higher gsw and E in *G. bickii* across non-control treatments (Fig. 4A, B).

Leaf structural plasticity differs between *G. hirsutum* and *G. bickii* under water deficit.

Leaf thickness was consistently greater (31 – 53%) in *G. hirsutum* compared to *G. bickii* across all treatments (Fig. 5A). A significant increase in leaf thickness was observed only under soil water deficit at 32°C, relative to well-watered conditions at the same temperature, where the degree of leaf thickening was greater for *G. hirsutum* (26%) than *G. bickii* (19%). Mesophyll thickness was strongly and positively correlated with total leaf thickness (*p* <0.001; *r* = 1.00; Table S1), accounting on average for ∼90% of leaf thickness.

**Figure 5.**
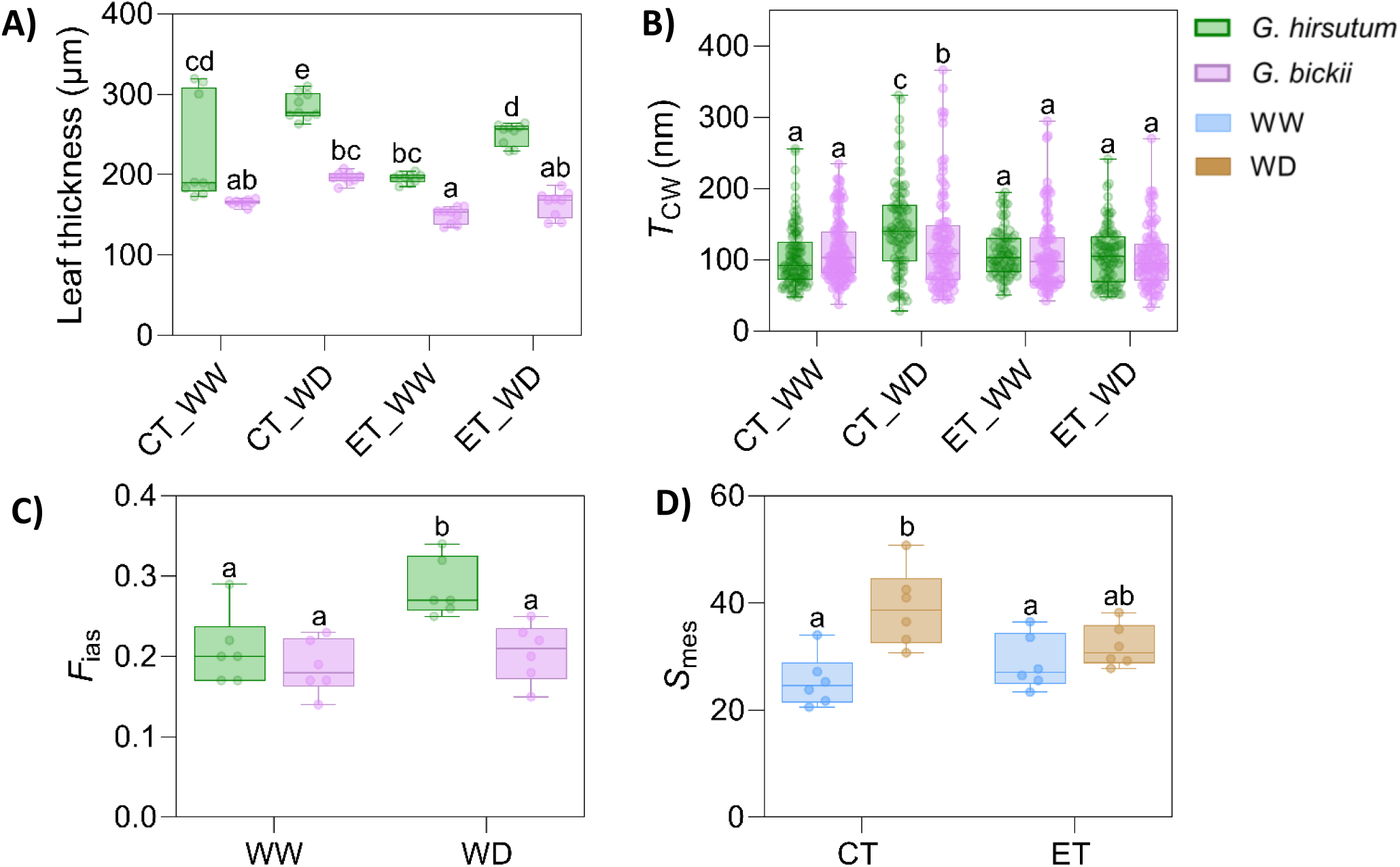
Leaf anatomical traits of *G. hirsutum* (green) and *G. bickii* (pink) under combinations of control (CT) and elevated (ET) temperatures and well-watered (WW) and water-deficit (WD) treatments. A) Leaf thickness (*n*=9). B) Mesophyll cell wall thickness (*T*CW; *n*=216). C) Fraction of mesophyll occupied by intercellular airspace (*F*ias; *n*=6) shown for WW and WD treatments. D) Mesophyll surface area exposed to intercellular airspace (*S*mes; *n*=6) shown across CT and ET with WW and WD treatments as coloured boxes. For panels A, and B, treatments are indicated as CT_WW, CT_WD, ET_WW, and ET_WD. Boxes indicate the interquartile range (25th–75th percentile), the horizontal line inside each box represents the median, and whiskers show the minimum and maximum values. Raw data points are overlaid (jittered for clarity). Different letters above boxes denote significant differences between treatments according to Fisher’s protected LSD (α = 0.05).

*T*CW was affected by water availability (*p* <0.001), temperature (*p* <0.001), and their interactions with species (*species × water, p* = 0.003; *water × temperature, p* <0.001; Fig. 5B). *T*CW was unchanged under elevated temperature (38°C) regardless of water status, compared to well-watered conditions at 32°C. However, under soil water deficit at 32°C, cell walls thickened in both species – more substantially in *G. hirsutum* (31% increase) than in *G. bickii* (16% increase). Under this treatment, *G. hirsutum* also exhibited significantly thicker cell walls than *G. bickii*.

Species and water availability interacted to influence the fraction of mesophyll occupied by intercellular airspace (*F*ias; *p* = 0.002; Fig. 5C). Under well-watered conditions, *F*ias was similar between species, but under water deficit, *F*ias increased in *G. hirsutum* (37% relative to well-watered) while remaining stable in *G. bickii*.

Water availability (*p* = 0.008) and temperature (*p* = 0.006) influenced the surface are of mesophyll cells exposed to the intercellular airspace (*S*mes; Fig. 5D). *S*mes remained relatively stable under well-watered conditions at both 32°C and 38°C, and was also unchanged under soil water deficit at 38°C. However, under soil water deficit at 32°C, *S*mes increased by 54% compared to well-watered conditions at the same temperature.

Pearson correlation analysis across species and treatments (Table S1) revealed that *g*m correlated most strongly with *A*/*g*m (*p* < 0.001; *r* = −0.86), *E* (*p* < 0.001; *r* = 0.69), *g*sw (*p* < 0.001; *r* = 0.66), *C*i (*p* < 0.001; *r* = 0.64), and *C*i– *C*a (*p* < 0.001; *r* = 0.64), followed by leaf and mesophyll thickness (*p* < 0.01; *r* = −0.56) and *S*mes (*p* < 0.05; *r* = −0.47). *T*CW showed a weaker, non-significant negative correlation with *g*m (*r* = −0.33, n.s.). Leaf thickness was also positively associated with *F*ias (*p* < 0.001; *r* = 0.64), *S*mes (*p* < 0.01; *r* = 0.55), and *A*/*g*m (*p* < 0.01; *r* = 0.52), and negatively with *g*m (*p* < 0.01; *r* = −0.56), *g*sw (*p* < 0.01; *r* = −0.61), *E* (*p* < 0.01; *r* = −0.52), and *C*i and *C*i– *C*a (*p* < 0.01; *r* = −0.62).

Significant correlations between *g*m and both leaf thickness and mesophyll thickness were observed across species, water availability, and temperature, with species showing the strongest relationships (*p* <0.001; *r* = 0.44; Table S1). Across treatments, *G. hirsutum* showed significant negative correlations between *g*m and leaf thickness (*p* = 0.0022; R^2^ = 0.625; Fig. 6A) and mesophyll thickness (*p* = 0.0019; R^2^ = 0.637; Fig. 6B), whereas no such relationships were observed in *G. bickii*. No significant relationship was observed between *g*m and *F*ias (Fig. 6C) in either species or treatment. A negative linear relationship was detected between *g*m and *S*mes under control temperatures (*p* = 0.0016; R^2^ = 0.647; Fig. 6D), but not under elevated temperature. Additionally, *g*m was negatively associated with *T*CW under water deficit conditions (*p* = 0.0167; R^2^ = 0.451; Fig. 6E), with no relationship detected under well-watered conditions.

**Figure 6.**
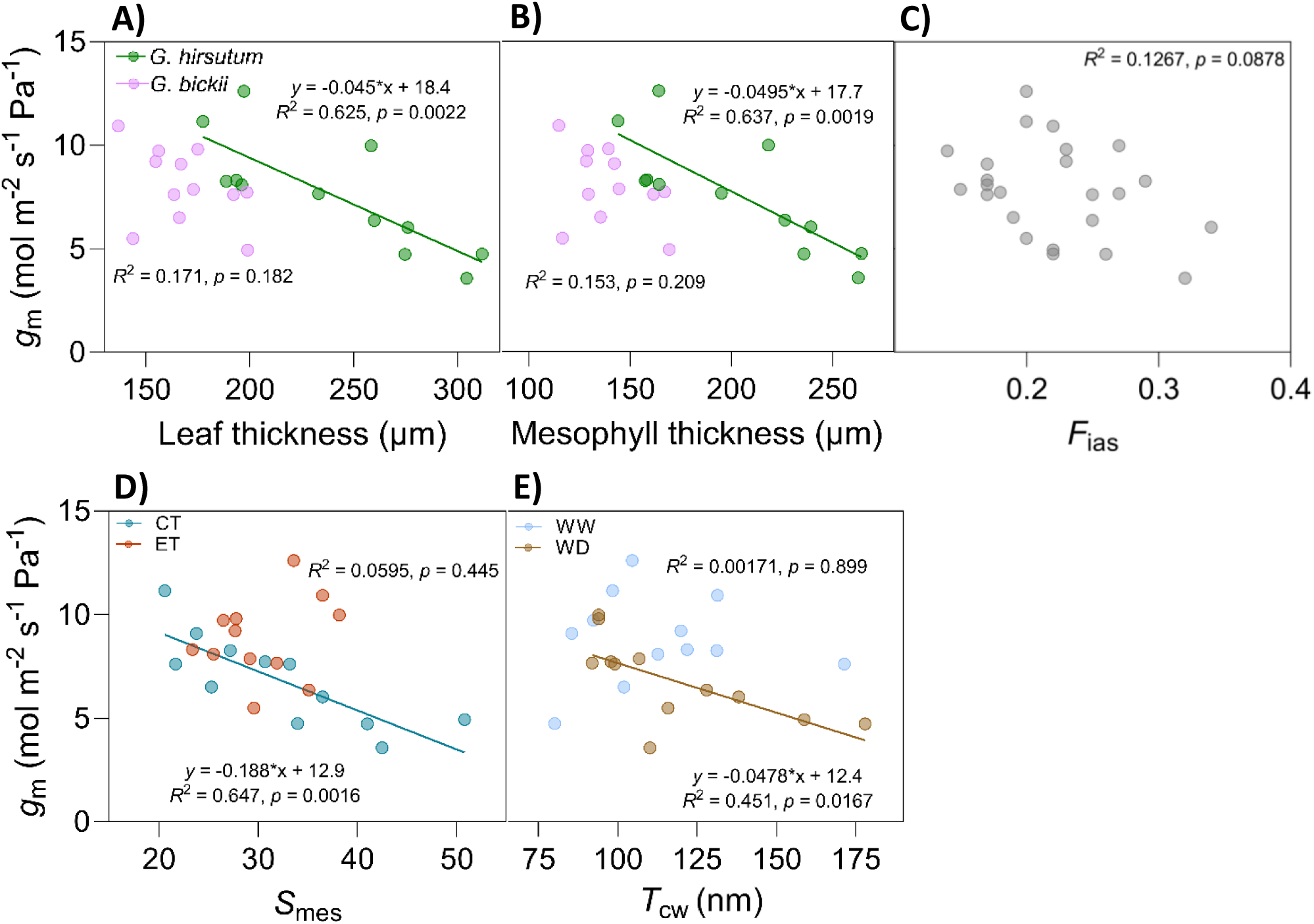
Regression analysis between mesophyll conductance (*g*m) and anatomical parameters A) leaf thickness (*n*=12), B) mesophyll thickness (*n*=12), C) fraction of mesophyll occupied by intercellular airspace (*F*ias; *n*=24), D) surface area of mesophyll cells exposed to intercellular airspace (*S*mes; *n*=12), and E) cell wall thickness (*T*CW; *n*=12). Group/treatment effects determined by ANCOVA/multiple linear regression analysis were detected for A), B), D), and E), represented by different coloured symbols for species (*G. hirsutum =* green; *G. bickii* = pink), water treatments (well-watered (WW) = blue; water-deficit (WD) = brown), and temperature treatments (control temperature (CT) = teal; elevated temperature (ET) = red). Separate regression lines represent significant group/treatment effects (*p*<0.05).

*A*380 was positively correlated with *g*m (*p* = 0.044; R^2^ = 0.171; Fig. 7A) and *g*sw (*p* < 0.01; *r* = 0.54; Table S1) and negatively correlated with *S*mes (*p* = 0.0022; R^2^ = 0.625; Fig. 7B; Table S1). *A*380 did not correlate with *C*i (*p* = n.s.; *r* = 0.34; Table S1), *A*/*g*m (*p* = n.s.; *r* = 0.002; Table S1) or *C*i-*C*a (*p* = n.s.; *r* = 0.34; Table S1).

**Figure 7.**
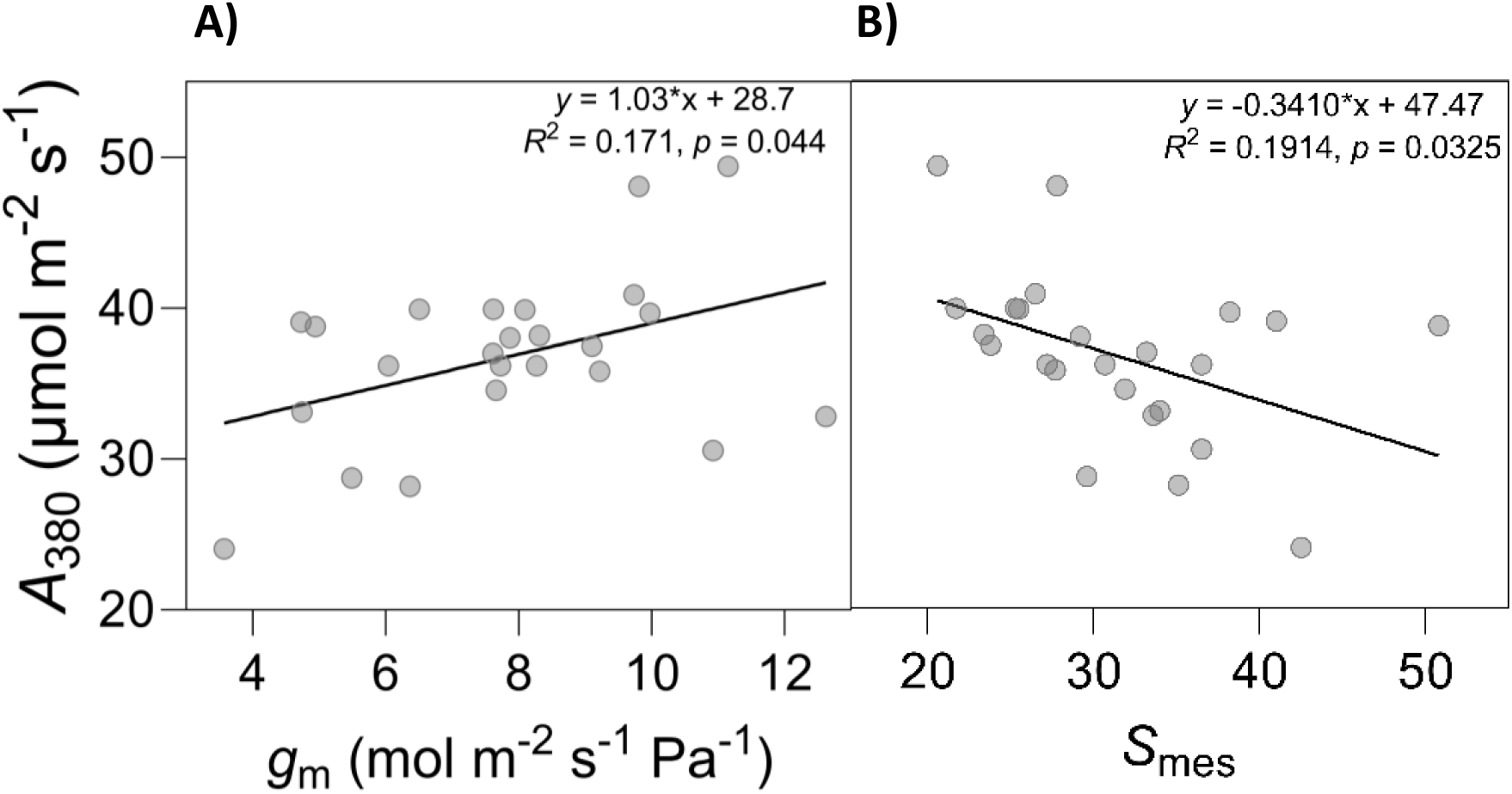
Regression analysis between net leaf photosynthesis at 380 µmol CO2 mol^-1^ air (*A*380) and A) mesophyll conductance (*g*m; *n*=24), and B) surface area of mesophyll cells exposed to intercellular airspace (*S*mes; *n*=24).

*F*ias exhibited significant positive relationships with both mesophyll thickness (*p* <0.001; *r* = 0.65; Table S1) and leaf thickness, with the relationship of leaf thickness being significant only under water-deficit conditions (*p* = 0.0015; R^2^ = 0.653; Fig. 8A). *S*mes was also positively associated with leaf thickness (*p* = 0.0053; R^2^ = 0.3031; Fig. 8B), mesophyll thickness (*p* <0.01; *r* = 0.58 Table S1) and *F*ias (*p* <0.01; *r* = 0.55; Table S1). *A*/*g*m was positively correlated (Table S1) with *T*CW (*p* <0.05; *r* = 0.40), *S*mes (*p* <0.05; *r* = 0.47), mesophyll thickness (*p* <0.05; *r* = 0.51) and leaf thickness (*p* <0.01; *r* = 0.52).

**Figure 8.**
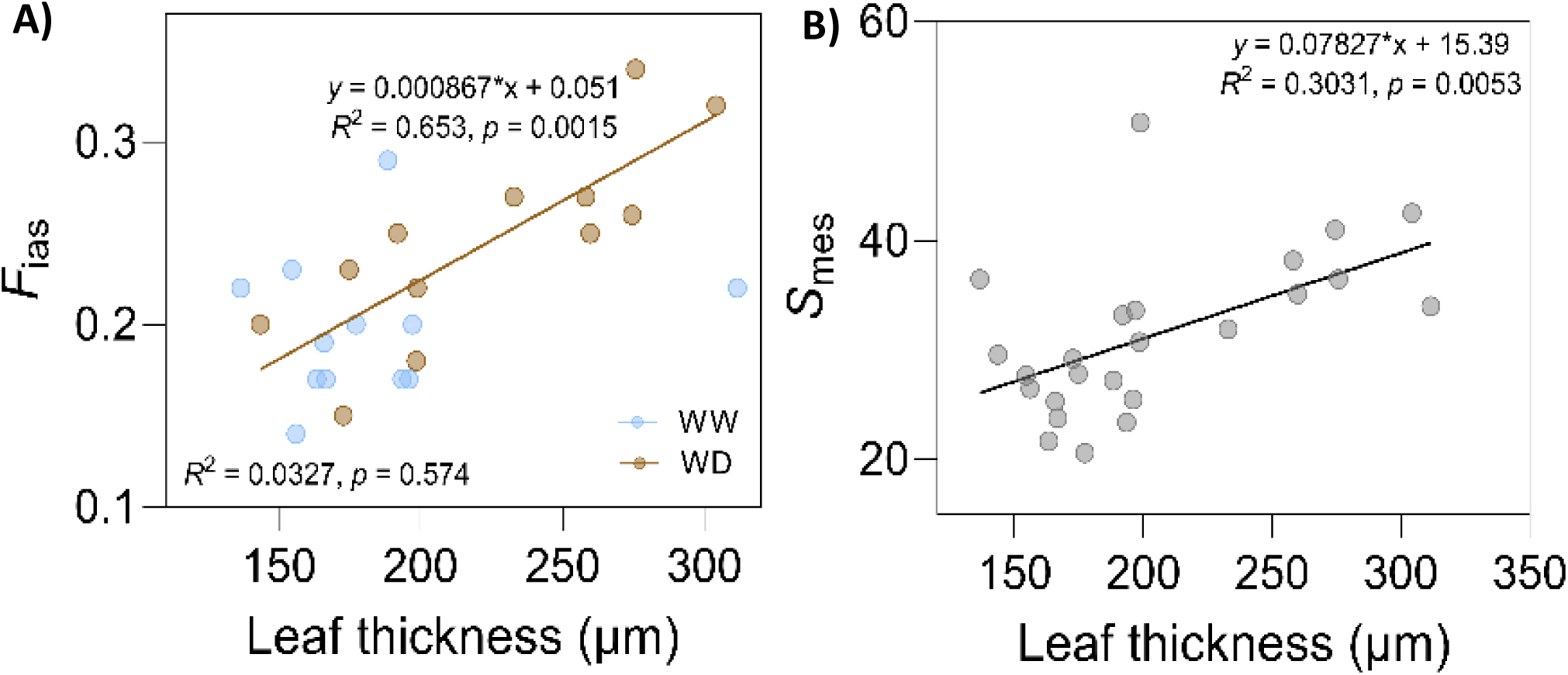
Relationships between leaf thickness and mesophyll structural traits. A) Fraction of mesophyll occupied by intercellular airspace (*F*ias) plotted against leaf thickness for well-watered (WW; blue points) and water-deficit (WD; brown points) leaves. A significant positive regression was observed for WD only (*p*=0.0015, R² = 0.653). B) Mesophyll surface area exposed to intercellular airspace (*S*mes) plotted against leaf thickness across all leaves (*p*=0.0053, R² = 0.3031). Raw data points are shown for transparency, with regression lines overlaid where significant.

## Discussion

Mesophyll conductance (*g*m) has emerged as a critical determinant of photosynthetic performance by governing the diffusion of CO2 from substomatal cavities to chloroplasts. Although widely recognized as finite and variable, the mechanistic basis of *g*m diversity, particularly under environmental stress, remains unclear (Evans, 2021, Flexas et al., 2008). Our study addresses this by characterising the temperature sensitivity of *g*m across diverse *Gossypium* species, followed by detailed examination of commercially cultivated cotton (*G. hirsutum*) and Australian wild species *G. bickii* grown under elevated temperature and soil water deficit. This two-phase design enables mechanistic insight into species-specific *g*m plasticity and its influence on photosynthetic performance under abiotic stress.

### Thermal sensitivity of mesophyll conductance underpins a trade-off between capacity and stability of photosynthesis across *Gossypium* species

*Gossypium* species differed in both the magnitude and thermal sensitivity of *g*m. Thermal optima of *g*m and photosynthesis occurred between 33 and 36°C, but responses above the optima diverged among species. *G. hirsutum* exhibited high *g*m and *A*380 near optimal temperatures, followed by a modest decline above the optima, whereas *G. bickii* maintained comparatively stable *g*m and photosynthetic rates across the temperature range examined (Fig. 1B). *G. gossypioides*, which had inherently low *g*m, showed minimal post-optimum decline in both *g*m and *A*380. Together, these contrasting patterns suggest a trade-off between photosynthetic capacity and thermal resilience, whereby *G. hirsutum* maximises assimilation under favourable conditions but exhibits greater sensitivity to heat, while some wild species adopt more conservative yet thermally stable strategies.

Short-term temperature responses of *g*m have been reported to follow three general patterns: (i) continuous increases with temperature; (ii) weak temperature sensitivity; and (iii) increases followed by declines at high temperature (Huang et al., 2022, Li et al., 2020, von Caemmerer and Evans, 2015). However, it remains unresolved whether these contrasting patterns primarily reflect biological variation among species or methodological challenges associated with estimating *g*m across wide temperature ranges, particularly at extreme temperatures.

Comparisons across studies indicate that both factors may contribute. For example, *G. hirsutum* measured using carbon isotope discrimination by von Caemmerer and Evans (2015) exhibited a continuous increase in *g*m with temperature, reaching values substantially higher than observed here. Although the same isotope-based approach was used in the present study, the modest reductions in *g*m observed at 40°C occurred near the upper limits of reliable estimation, suggesting that uncertainty in *g*m calculations may increase at extreme temperatures. Differences in growth conditions may also contribute, as *G. hirsutum* plants in von Caemmerer and Evans (2015) were grown under cooler conditions (28/18°C day/night) than in the present study, highlighting the potential importance of thermal acclimation.

Declines in *g*m at high temperature have been reported previously using fluorescence-based methods (Bernacchi et al., 2002) and, in some cases, isotope discrimination (Yamori et al., 2006); however, these responses are not consistently observed across species or measurement techniques. Indeed, isotope-based studies in tobacco have not detected comparable declines (von Caemmerer and Evans, 2015), suggesting that apparent reductions in *g*m at high temperature may, in some instances, arise from methodological artefacts rather than reflecting intrinsic physiological limitations. Accordingly, declines in *g*m at supra-optimal temperatures should be interpreted cautiously.

The temperature sensitivity of *g*m is commonly interpreted using a two-component diffusion model that partitions *g*m into liquid-phase (*g*liq) and membrane-phase (*g*mem) components (Evans and von Caemmerer, 2013, von Caemmerer and Evans, 2015). *g*liq, encompassing diffusion through cell walls, cytoplasm, and chloroplast stroma, is primarily determined by the physical properties of water and the effective CO2 diffusion path length, and is expected to show relatively weak temperature sensitivity (Li et al., 2020, von Caemmerer and Evans, 2015, Walker et al., 2013). In contrast, *g*mem, reflecting CO2 permeability across the plasma membrane and chloroplast envelope and regulated in part by aquaporins, is highly temperature-dependent and expected to increase exponentially with temperature (Li et al., 2020, von Caemmerer and Evans, 2015). While this framework provides a useful basis for interpreting variation in *g*m temperature sensitivity, it cannot simulate the decline in *g*m observed at supra-optimal temperatures (Li et al., 2020), and empirical estimates of the activation energy governing membrane CO2 permeability remain unavailable (von Caemmerer and Evans, 2015). Consequently, this framework is better suited to interpreting variation in the thermal stability of *g*m than to mechanistically resolving declines at high temperature, which may involve additional processes such as heat-induced constraints on membrane permeability or aquaporin function.

Analyses of *A*/*g*m and *A*/*g*sc further revealed that the relative contributions of stomatal and mesophyll limitations to photosynthesis differed among species and shifted with temperature. At high temperatures, *G. arboreum*, *G. gossypioides* and *G. hirsutum* exhibited the largest *C*a-*C*i drawdowns, indicating increasing stomatal limitation. In contrast, *G. bickii* maintained low and thermally stable *A*/*g*sc, consistent with its higher *g*sw and *C*i (Table S2). In *G. hirsutum*, *A*/*g*m increased above the thermal optimum, indicating that mesophyll limitation became increasingly dominant as *g*m declined more strongly with temperature. By contrast, *G. bickii* maintained low and stable *A*/*g*sc and *A*/*g*m across temperatures, reflecting coordinated diffusional stability through both stomatal and mesophyll pathways. This diffusional robustness likely underpins its sustained photosynthetic performance at temperatures approaching 40°C, highlighting *g*m stability rather than maximal capacity as a key determinant of thermal tolerance among *Gossypium* species.

### Contrasting anatomical strategies underpin species-specific mesophyll conductance responses to water deficit

Soil water deficit reduced *g*m in both *G. hirsutum* and *G. bickii* at 32°C, with a greater decline in *G. hirsutum*, indicating greater sensitivity to water limitation (Fig. 3A). *A*380 remained stable under individual stresses but declined substantially in *G. hirsutum* under combined elevated temperature and soil water deficit (Fig. 3B). While reductions in *g*sw contributed to decreased assimilation (Table S2), the pronounced decline in *g*m highlights its central role in constraining photosynthesis under combined stress. *g*m, which integrates CO₂ diffusion from the intercellular airspace to the chloroplast stroma, is strongly influenced by species-specific anatomical traits. Our results reveal contrasting strategies underpinning *g*m plasticity between the two species.

*Mesophyll restructuring in G. hirsutum under water stress limits mesophyll conductance G. hirsutum* had inherently thicker leaves than *G. bickii*, consistent with previous comparisons between domestic cotton against wild genotypes (Lei et al., 2022a). Under soil water deficit, *G. hirsutum* exhibited further leaf and mesophyll thickening, similar to previous reports for cultivated cotton (Han et al., 2019). Notably, this thickening occurred alongside pronounced mesophyll reconfiguration, characterised by increased leaf porosity (*F*ias) and mesophyll surface exposure to the intercellular airspace (*S*mes), indicating a loosening of internal leaf structure.

Despite these structural adjustments, neither *g*m nor *A*380 increased under water deficit, suggesting that expansion of diffusive surface area alone was insufficient to sustain CO2 supply to the chloroplast. This is consistent with the minor contribution of gas-phase conductance (*g*ias), which depends on leaf thickness, *F*ias, and the tortuosity of diffusion pathways (Hanba et al., 1999, Syvertsen et al., 1995), relative to the much larger resistance imposed by liquid-phase diffusion, where CO2 diffusion is ∼10,000-fold slower than in air (Evans et al., 1994, Terashima et al., 2006). Consequently, increases in *F*ias and *S*mes in *G. hirsutum* may represent compensatory or multifunctional responses to water deficit, potentially conferring non-diffusional benefits such as enhanced convective cooling (Lombardini and Rossi, 2019), rather than alleviating mesophyll diffusion limitation (Flexas et al., 2008, Flexas et al., 2009, Genty et al., 1998, Piel et al., 2002, Terashima et al., 2006, Terashima et al., 2011).

Across species, *S*mes was negatively correlated with *g*m under control temperature and with *A*380. This contrasts with the generally positive association between *S*mes and *g*m reported across many herbaceous and woody species (Flexas et al., 2008, Kogami et al., 2001, Lloyd et al., 1992, Miyazawa and Terashima, 2001, Ouyang et al., 2017, Syvertsen et al., 1995, Terashima et al., 2006). However, decoupled and negative relationships between *S*mes and *g*m have been reported previously in deciduous tree species (Hanba et al., 2001) and in transgenic rice overexpressing the barley aquaporin HvPIP2;1 (Hanba et al., 2004), respectively. In the present study, this negative relationship suggests that greater *S*mes did not improve CO2 diffusion because downstream liquid-phase resistances may have dominated *g*m responses.

Notably, cytoplasmic diffusion and *S*c are known to strongly influence *g*m and photosynthesis (e.g., (Ouyang et al., 2017, Tholen et al., 2008, Tosens et al., 2012a, Zhang et al., 2015, Zou et al., 2022)). *S*c was not quantified in this study, as chloroplast distribution across mesophyll cells showed pronounced heterogeneity. Resolving this variability would have required substantially greater sampling intensity than was feasible within the comparative framework of this work. However, domesticated cotton genotypes have previously been shown to exhibit lower cytoplasmic conductance (*g*cyt) than wild relatives, attributed to increased separation between the plasmalemma and chloroplast envelope (Lei et al., 2022a). Chloroplast migration away from the cell wall under water deficit stress could further lengthen the cytoplasmic diffusion pathway (Zhang et al., 2015). Such anatomical constraints could explain why increases in *S*mes were insufficient to prevent declines in *g*m under water deficit in *G. hirsutum*.

By contrast, *G. bickii* maintained thinner leaves that did not exhibit water deficit-induced thickening, while increasing *S*mes without altering *F*ias. This pattern is consistent with mesophyll reorganisation into smaller, more numerous cells (Fig. S4). This more targeted adjustment may enhance liquid-phase diffusion efficiency at the chloroplast interface without the broader structural trade-offs observed in *G. hirsutum*. This strategy likely contributed to the smaller reduction in *g*m under water deficit.

#### Cell wall reinforcement in G. hirsutum amplifies diffusional resistance under water deficit

In addition to mesophyll reorganisation, cell wall thickening under water deficit imposed further resistance to CO2 diffusion. Cell wall reinforcement was more pronounced in *G. hirsutum* than in *G. bickii* under soil water deficit at 32°C, consistent with a common drought response across species (Giuliani et al., 2013, Ouyang et al., 2017, Scafaro et al., 2011, Tosens et al., 2012a). Accordingly, *g*m was inversely related to *T*CW under water deficit (Fig. 6E), consistent with previous findings that thicker cell walls impede CO₂ diffusion and can contribute >25% to total mesophyll resistance (Evans et al., 2009, Evans and von Caemmerer, 2013, Flexas et al., 2021, Galmés et al., 2013, Han et al., 2016, Huang et al., 2022, Ouyang et al., 2017, Salesse-Smith and Xiao, 2025, Scafaro et al., 2011, Terashima et al., 2011, Tholen and Zhu, 2011, Tosens et al., 2012a, von Caemmerer and Evans, 2015). This also reinforces the conclusion from the previous subsection that increased *S*mes was insufficient to overcome resistance imposed by increased liquid-phase diffusion distances such as thicker cell walls.

The smaller reduction in *g*m under water limitation in *G. bickii* is consistent with its more conservative cell wall thickening. However, variation in *g*m is unlikely to be explained by *T*CW alone, particularly when *T*CW is below ∼0.4µm, as observed here (Lei et al., 2022a). Instead, multiple studies highlight the importance of liquid-phase properties downstream of wall thickness, including stromal conductance associated with chloroplast width, as well as cell wall effective porosity and composition, in driving *g*m variation (Carriquí et al., 2019, Carriquí et al., 2020, Han et al., 2019, Huang et al., 2022, Pathare et al., 2024, Salesse-Smith et al., 2024, Salesse-Smith and Xiao, 2025, Tosens et al., 2012a, Tosens et al., 2016). Consistent with this, wild *Gossypium* genotypes possess pectin-rich cell walls with low crystallinity (Sun et al., 2025b), forming a more flexible matrix that facilitates liquid-phase CO2 diffusion (Flexas et al., 2021). Such wall properties likely contribute to the greater stability of *g*m observed here in *G. bickii* under elevated temperature and soil water deficit.

#### Biochemical and biophysical factors may modulate mesophyll conductance independently of anatomy

Although anatomical traits explained a substantial fraction of *g*m variation across species and treatments, anatomy alone cannot fully account for the observed responses. This is particularly evident from i) the decoupling between anatomical traits and *g*m under specific treatments, and ii) the stimulatory effect of elevated temperature on *g*m in the absence of measurable anatomical adjustment. These patterns indicate that biochemical and biophysical processes contribute to *g*m regulation alongside structural determinants.

Experimental and modelling studies demonstrate that CO2 diffusion across membranes is dynamically regulated by aquaporins, carbonic anhydrase activity, and membrane fluidity, which modulate *g*m independently of anatomy or interact with structural traits to shape temperature and stress responses (Bernacchi et al., 2002, Evans, 2021, Flexas et al., 2008, Han et al., 2016, Han et al., 2019). In particular, aquaporin-mediated CO2 permeability exhibits strong temperature sensitivity and has been implicated in rapid changes in *g*m at high temperatures (Bernacchi et al., 2002, Hanba et al., 2004, Scafaro et al., 2011, Uehlein et al., 2008).

### Heat stress exacerbates water deficit by limiting CO2 diffusion plasticity

Despite significant shifts in *g*m with water deficit and elevated temperature applied individually, *A*380 remained buffered for both *G. hirsutum* and *G. bickii*, indicating that photosynthesis was CO2 diffusion was not the sole limitation under single stress conditions. However, when heat and water deficit co-occurred, this buffering capacity collapsed in *G. hirsutum*, revealing a critical interaction between thermal and hydraulic stress.

Elevated temperature constrained the structural plasticity typically induced by water deficit, as leaf and mesophyll thickening, *S*mes and *T*CW became unresponsive to water deficit at 38°C. This suppression of anatomical acclimation suggests that heat stress imposes a distinct constraint on anatomical plasticity. Under well-watered conditions, elevated temperature increased *g*m without concomitant anatomical adjustments, consistent with enhanced membrane fluidity and permeability, and aquaporin activity (Bernacchi et al., 2002, Evans, 2021, Flexas et al., 2006b, Hanba et al., 2004, Scafaro et al., 2011). Under combined heat and water deficit, however, this temperature-induced stimulation of *g*m was negated, consistent with impaired aquaporin function (Bernacchi et al., 2002), leading to reduced CO2 assimilation in *G. hirsutum*.

Together, these findings indicate that heat stress exacerbates water stress limitation by constraining both structural and biochemical pathways that normally sustain CO2 diffusion. *G. hirsutum*, which relies more heavily on structural acclimation to maintain *g*m, therefore exhibits a greater mismatch between CO2 supply and biochemical demand under the combined stress. By contrast, *G. bickii* maintains relatively stable *g*m and *A*380, potentially due to superior cell wall properties (Sun et al., 2025b) and greater reliance on liquid-phase diffusion efficiency, conferring resilience to concurrent heat and water deficit stress.

## Conclusion

Our findings identify mesophyll conductance (*g*m) as both a vulnerability in *G. hirsutum* and a key determinant of photosynthetic responses to abiotic stress across *Gossypium* species. Species differed markedly in both the magnitude and thermal stability of *g*m, revealing a trade-off between high photosynthetic capacity near optimal temperatures and resilience under supra-optimal conditions. Under water deficit, cultivated *G. hirsutum* exhibited substantial anatomical reorganisation, including increased leaf porosity, cell wall thickness and mesophyll surface exposure to the intercellular airspace, yet these adjustments did not prevent declines in *g*m, indicating that downstream liquid-phase resistances dominate diffusion limitation under stress. By contrast, the Australian native species *G. bickii* maintained comparatively stable *g*m across heat and water stress, consistent with more conservative and targeted adjustments that support efficient liquid-phase diffusion, without broad structural modifications, providing a more effective response to stress that maintains CO₂ diffusion.

These divergent responses demonstrate that *g*m is not determined by static leaf anatomy alone, but by the dynamic coordination of anatomical, biophysical, and biochemical properties. Importantly, our results show that CO2 supply bottlenecks are dynamic: stomatal limitations increase in some species at supra-optimal temperatures, while mesophyll limitations either remain relatively stable or vary according to *g*m plasticity. This dynamic behaviour highlights the need to explicitly account for mesophyll conductance in mechanistic models of photosynthesis, as treating *g*m as infinite or static, or neglecting species-specific differences can lead to substantial errors in the estimation of biochemical parameters such as *V*cmax, particularly under climatic extremes (Bernacchi et al., 2002, Flexas et al., 2008, Sargent et al., 2024).

By integrating cross-species comparisons with anatomical and physiological analyses, this study refines mechanistic understanding of *g*m regulation and plasticity under extreme environments, and provides a framework for exploring how anatomical, biophysical, and biochemical diversity underpin photosynthetic resilience. Future research combining physiology, omics, biochemistry, and modelling will be critical for quantifying these interacting components and for informing strategies to enhance *g*m plasticity and climate resilience in both agricultural and ecological systems.

## Supporting information

Supplementary Data_Raw Temperature Response Data

Supplementary Data_Raw Water Deficit x Elevated Temperature Experiment Data

Supplementary Data_Raw Microscopy Data

## Acknowledgements

This work was supported by the Cotton Research and Development Corporation (project number UWS2201 and ABARES Science and Innovation Award), the Australian Government through the Australian Research Council Centre of Excellence for Translational Photosynthesis (CE140100015), and CSIRO. Technical expertise and assistance provided by Dr. Daniel Fanna, Mr. Hyunsung Min and Ms. Sarah Fletcher (AMCF), Ms. Sue Lindsay and Dr. Chao Shen (MAFF), Dr. Rosemary White and Dr. Florence Danila (ANU), and Jiwon Lee (Centre for Advanced Microscopy, ANU) is gratefully acknowledged. RES was supported by the Western Sydney University VC Fellowship.

## Author contributions

D.S., R.S., and W.C. conceived the project and designed the experiments. J.E., and S.v.C. contributed to experimental design and data analysis. D.S. and G.D. implemented the water deficit treatments and collected gas exchange and leaf water potential data. D.S. and K.C. collected and analysed mesophyll conductance data. D.S., L.G., and S.L. processed and imaged microscopy samples. D.S. analysed all datasets. All authors contributed to writing the manuscript.

## Data availability

All data supporting the findings of this study are available within the supplementary materials and the Source data files published online.

## Supplemental Data

**Table S1.** Pearson’s correlation matrix (lower triangle) showing relationships between key gas exchange and anatomical parameters measured from the elevated temperature by water deficit experiment. Data were pooled across all species and treatments within this experiment. Values represent Pearson’s correlation coefficients (*r*). Statistical significance is indicated by asterisks: *p*<0.05 (\**), p*<0.01 *(**), and p*<0.001 *(\*\*\**). Sample size varied among parameter pairs due to differences in measurement techniques

| | $A/g_m$ | $C_i$ | $C_i-C_a$ | $g_{sw}$ | $g_m$ | $A_{380}$ | $E$ | $VPD_L$ | $T_{CW}$ | $F_{ias}$ | $S_{mes}$ | Meso.<br>Thick. |
| --- | --- | --- | --- | --- | --- | --- | --- | --- | --- | --- | --- | --- |
| $A/g_m$ | | | | | | | | | | | | |
| $C_i$ | -0.49* | | | | | | | | | | | |
| $C_i-C_a$ | -0.49* | 1.00*** | | | | | | | | | | |
| $g_{sw}$ | -0.46* | 0.94*** | 0.94*** | | | | | | | | | |
| $g_m$ | -0.86*** | 0.64*** | 0.64*** | 0.66*** | | | | | | | | |
| $A_{380}$ | 0.002 | 0.34 | 0.34 | 0.54** | 0.41* | | | | | | | |
| $E$ | -0.61** | 0.64*** | 0.64*** | 0.73*** | 0.69*** | 0.38 | | | | | | |
| $VPD_L$ | 0.003 | -0.76*** | -0.77*** | -0.72*** | -0.30 | -0.52** | -0.13 | | | | | |
| $T_{CW}$ | 0.40* | -0.14 | -0.14 | -0.17 | -0.33 | -0.07 | -0.29 | -0.09 | | | | |
| $F_{ias}$ | 0.26 | -0.48* | -0.48* | -0.54** | -0.36 | -0.34 | -0.54** | 0.25 | 0.19 | | | |
| $S_{mes}$ | 0.47* | -0.52** | -0.52** | -0.61** | -0.47* | -0.44* | -0.53** | 0.29 | 0.32 | 0.55** | | |
| Meso. thick. | 0.51* | -0.62** | -0.62** | -0.62** | -0.56** | -0.36 | -0.53** | 0.36 | 0.04 | 0.65*** | 0.58** |  |
| Leaf thick. | 0.52** | -0.62** | -0.62** | -0.61** | -0.56** | -0.33 | -0.52** | 0.35 | 0.03 | 0.64*** | 0.55** | 1.00*** |

**Table S2.** Summary statistics and ANOVA results for key physiological traits measured in the temperature response experiment. Values are presented as means (± SD, *n*=5).

| Temp.<br>(°C) | Species | $g_m$ (mol m<sup>-2</sup> s<sup>-1</sup> Pa<sup>-1</sup>) | $A_{380}$ (μmol m<sup>-2</sup> s<sup>-1</sup>) | $g_{sw}$ (mol m<sup>-2</sup> s<sup>-1</sup>) | $C_i$ (μmol mol<sup>-1</sup>) | $E$ (mmol m<sup>-2</sup> s<sup>-1</sup>) | VPD<sub>L</sub> (kPa) | $A/g_m$ (Pa) | $A/g_{sc}$ (Pa) | $T_{leaf}$ (°C) |
| --- | --- | --- | --- | --- | --- | --- | --- | --- | --- | --- |
| 25 | *G. hirsutum* | 4.13 ± 0.68 | 30.65 ± 5.79 | 0.63 ± 0.19 | 276.80 ± 16.82 | 6.43 ± 0.92 | 1.20 ± 0.21 | 7.50 ± 1.32 | 8.23 ± 1.74 | 26.92 ± 0.45 |
|  | *G. arboreum* | 5.16 ± 1.07 | 30.74 ± 5.50 | 0.78 ± 0.26 | 287.80 ± 28.52 | 7.34 ± 1.16 | 1.17 ± 0.24 | 6.10 ± 1.41 | 7.13 ± 2.78 | 26.98 ± 0.43 |
|  | *G. bickii* | 4.25 ± 0.92 | 29.42 ± 6.47 | 1.17 ± 0.59 | 311.60 ± 19.90 | 7.17 ± 0.98 | 0.91 ± 0.23 | 7.17 ± 1.83 | 4.93 ± 1.95 | 26.03 ± 0.81 |
|  | *G. gossypoides* | 1.14 ± 0.55 | 9.45 ± 1.66 | 0.16 ± 0.05 | 262.70 ± 16.54 | 3.11 ± 0.64 | 2.00 ± 0.20 | 9.54 ± 3.31 | 10.25 ± 1.65 | 26.49 ± 0.34 |
|  | *G. sturtianum* | 4.01 ± 0.91 | 29.68 ± 2.31 | 0.72 ± 0.34 | 277.80 ± 39.62 | 6.45 ± 0.98 | 1.17 ± 0.30 | 7.64 ± 1.45 | 8.21 ± 3.88 | 26.64 ± 0.59 |
|  | ANOVA | <0.001 | <0.001 | <0.001 | <0.001 | <0.001 | <0.001 | <0.001 | <0.001 | <0.001 |
| 30 | *G. hirsutum* | 5.32 ± 0.93 | 39.09 ± 5.58 | 0.90 ± 0.22 | 281.80 ± 12.05 | 10.63 ± 1.84 | 1.43 ± 0.17 | 7.51 ± 1.39 | 7.20 ± 1.19 | 31.25 ± 1.18 |
|  | *G. arboreum* | 5.38 ± 1.12 | 38.60 ± 3.38 | 0.97 ± 0.24 | 286.80 ± 17.20 | 10.26 ± 1.40 | 1.32 ± 0.18 | 7.48 ± 1.41 | 6.78 ± 1.65 | 30.97 ± 0.98 |
|  | *G. bickii* | 4.78 ± 1.00 | 30.86 ± 3.51 | 1.14 ± 0.44 | 309.10 ± 17.19 | 10.25 ± 1.65 | 1.22 ± 0.18 | 6.74 ± 1.54 | 4.92 ± 1.65 | 30.54 ± 0.77 |
|  | *G. gossypoides* | 2.75 ± 0.64 | 18.01 ± 2.25 | 0.35 ± 0.11 | 270.20 ± 20.80 | 6.95 ± 1.46 | 2.13 ± 0.24 | 6.78 ± 1.20 | 9.00 ± 2.06 | 31.58 ± 1.07 |
|  | *G. sturtianum* | 5.29 ± 1.14 | 38.10 ± 4.47 | 1.25 ± 0.43 | 299.30 ± 23.14 | 11.23 ± 2.22 | 1.24 ± 0.19 | 7.49 ± 1.57 | 5.56 ± 2.37 | 31.04 ± 1.17 |
|  | ANOVA | <0.001 | <0.001 | <0.001 | <0.001 | <0.001 | <0.001 | 0.007 | <0.001 | <0.001 |
| 35 | *G. hirsutum* | 5.18 ± 0.66 | 41.28 ± 4.93 | 1.17 ± 0.24 | 293.40 ± 10.50 | 13.38 ± 2.04 | 1.48 ± 0.16 | 8.05 ± 1.07 | 5.87 ± 1.01 | 34.67 ± 1.29 |
|  | *G. arboreum* | 5.26 ± 0.80 | 37.15 ± 3.37 | 0.64 ± 0.16 | 254.20 ± 21.83 | 10.74 ± 1.37 | 1.92 ± 0.28 | 7.25 ± 1.41 | 9.85 ± 2.13 | 34.34 ± 1.19 |
|  | *G. bickii* | 5.49 ± 0.72 | 36.87 ± 4.73 | 1.35 ± 0.26 | 306.60 ± 14.34 | 12.50 ± 0.74 | 1.31 ± 0.14 | 6.81 ± 1.19 | 4.83 ± 1.22 | 33.73 ± 0.64 |
|  | *G. gossypoides* | 2.87 ± 0.62 | 19.63 ± 2.54 | 0.33 ± 0.11 | 254.10 ± 21.97 | 8.10 ± 1.36 | 2.62 ± 0.40 | 7.07 ± 1.40 | 10.30 ± 2.16 | 34.34 ± 0.72 |
|  | *G. sturtianum* | 5.29 ± 0.74 | 38.99 ± 2.23 | 1.03 ± 0.29 | 287.30 ± 17.14 | 12.00 ± 1.93 | 1.50 ± 0.25 | 7.55 ± 1.37 | 6.60 ± 1.69 | 33.92 ± 1.76 |
|  | ANOVA | <0.001 | <0.001 | <0.001 | <0.001 | <0.001 | <0.001 | <0.001 | <0.001 | 0.002 |
| 40 | *G. hirsutum* | 4.72 ± 1.11 | 36.19 ± 4.66 | 1.10 ± 0.58 | 280.80 ± 42.37 | 15.96 ± 3.60 | 2.05 ± 0.45 | 8.05 ± 2.35 | 7.07 ± 4.23 | 38.80 ± 1.74 |
|  | *G. arboreum* | 4.46 ± 1.60 | 28.48 ± 4.74 | 0.38 ± 0.25 | 226.70 ± 31.10 | 13.22 ± 2.58 | 3.54 ± 0.73 | 6.37 ± 1.25 | 12.20 ± 3.07 | 40.81 ± 0.53 |
|  | *G. bickii* | 5.13 ± 0.89 | 35.47 ± 3.79 | 1.33 ± 0.65 | 302.00 ± 18.53 | 17.86 ± 1.80 | 1.91 ± 0.34 | 7.08 ± 1.09 | 5.00 ± 1.68 | 39.57 ± 0.50 |
|  | *G. gossypoides* | 3.06 ± 0.93 | 19.07 ± 1.79 | 0.46 ± 0.20 | 270.90 ± 40.63 | 12.79 ± 3.82 | 3.10 ± 0.39 | 6.93 ± 2.45 | 8.31 ± 4.10 | 39.39 ± 1.79 |
|  | *G. sturtianum* | 4.95 ± 0.83 | 35.80 ± 2.16 | 1.19 ± 0.55 | 289.50 ± 37.55 | 16.91 ± 3.57 | 1.97 ± 0.33 | 7.17 ± 1.44 | 6.25 ± 3.65 | 39.18 ± 1.87 |
|  | ANOVA | <0.001 | <0.001 | <0.001 | <0.001 | <0.001 | <0.001 | <0.001 | <0.001 | <0.001 |

**Table S3.**
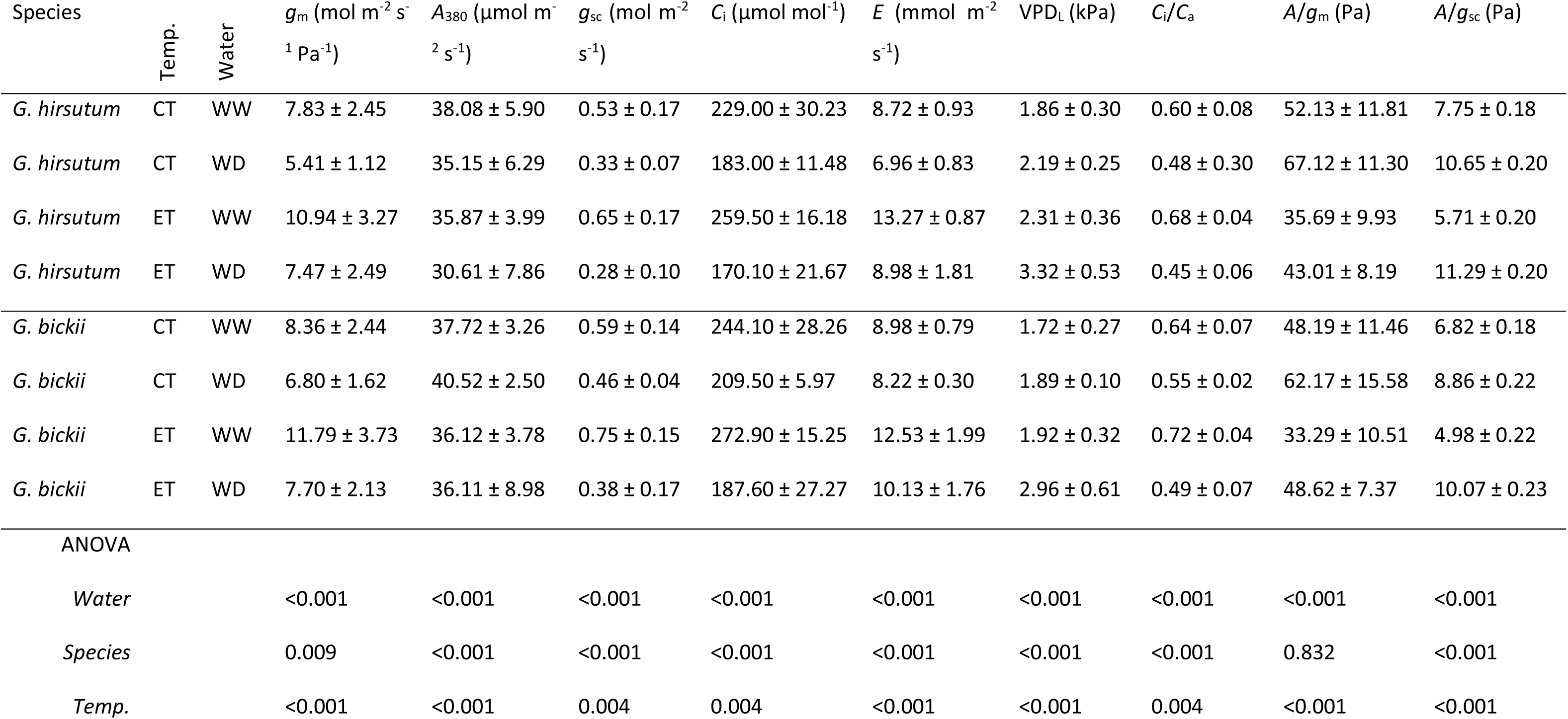
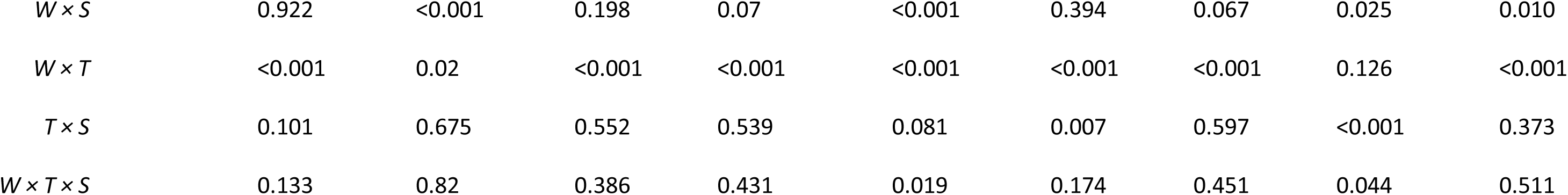
Summary statistics and ANOVA results for key physiological traits measured in the water-deficit × elevated-temperature experiment. CT = 32°C; ET = 38°C; WW = well-watered; WD = water-deficit. Values are presented as means of 60 technical measurements across six biological replicates. Error terms are ± SD for all variables except *A*/*g*sc, which is presented as ± SEM.

**Figure S1.**
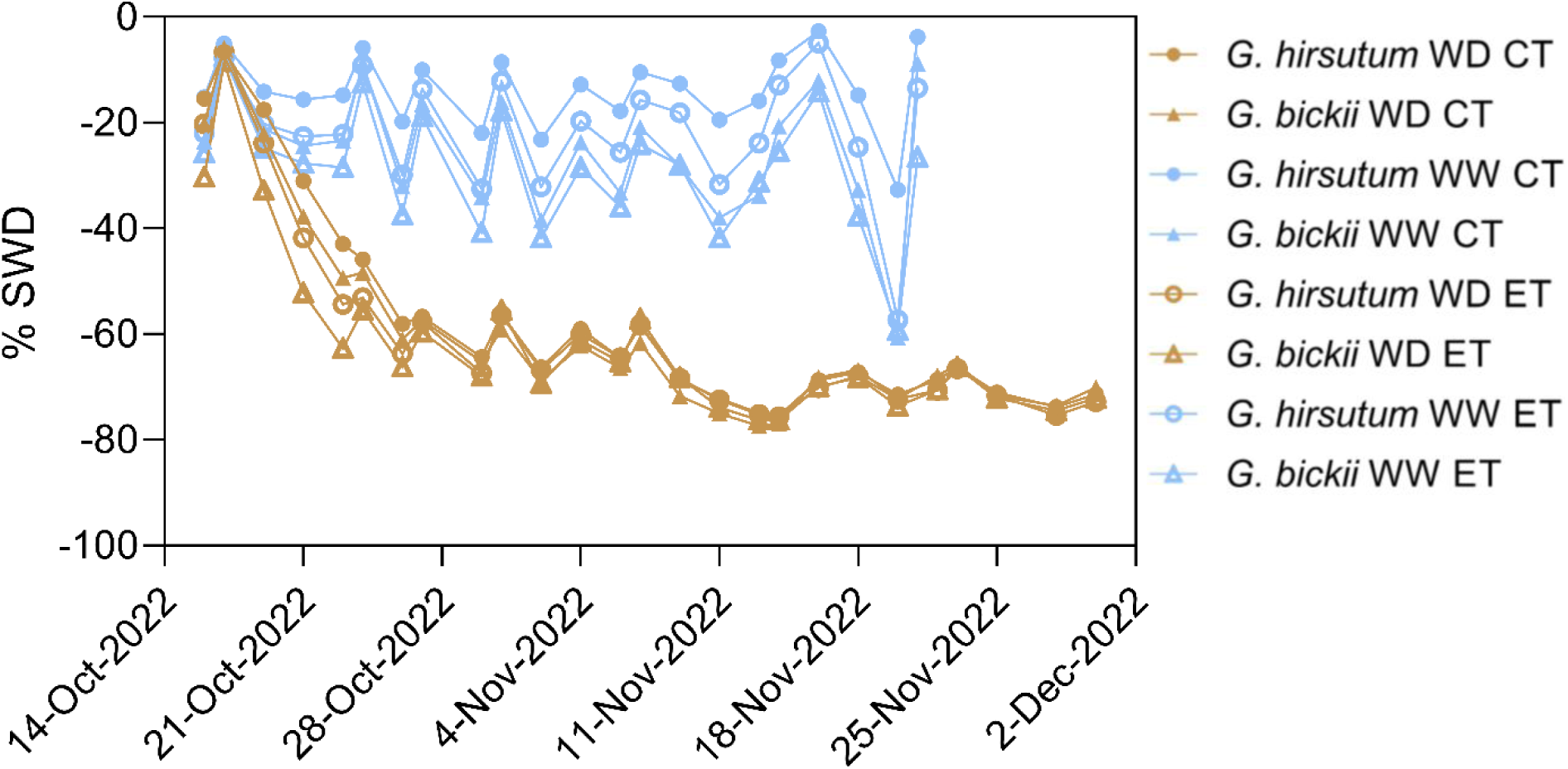
The pre-watering percentage soil water deficit (SWD) measured gravimetrically from 16^th^ October 2022. Water was withheld from water deficit-treated pots from 17^th^ October 2022, and soil water content was maintained at 60-75% SWD until experiment completion on 30^th^ November 2022.

**Figure S2.**
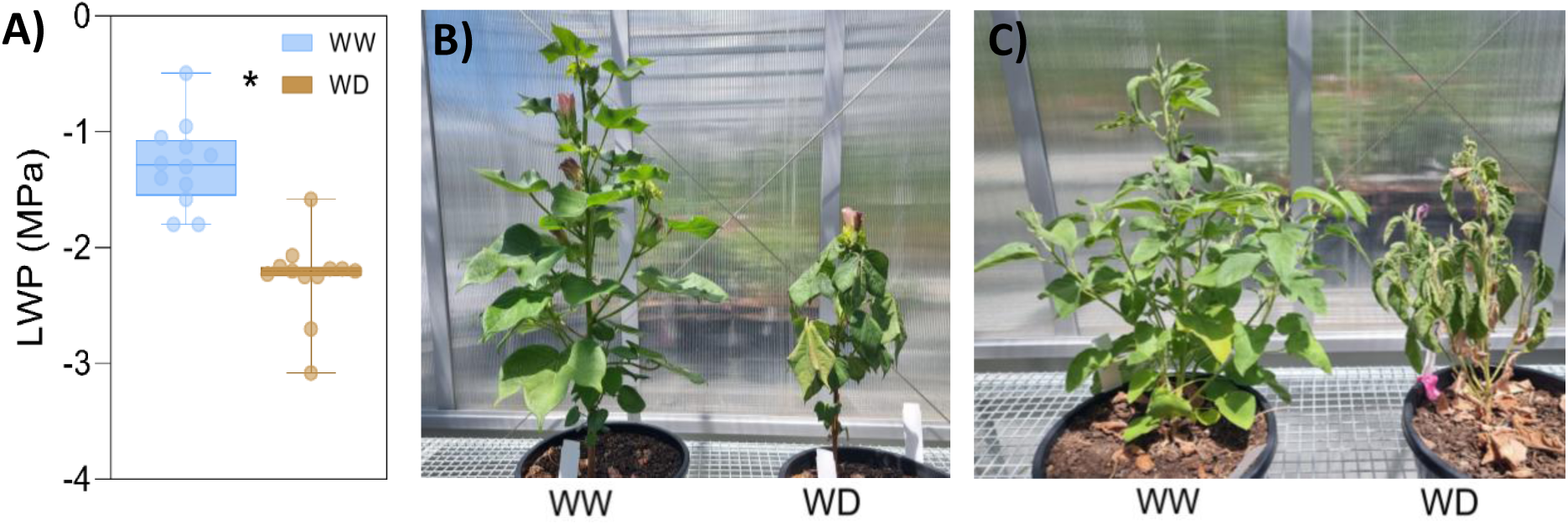
A) Pre-dawn leaf water potential (LWP) measured on 2^nd^ November 2022 on the first fully expanded leaf of the canopy of well-watered (WW; blue) and water-deficit (WD; brown) plants (*p* = 0.03, *n*=12) using a pressure chamber. Representative well-watered (left) and water-deficit (right) plants of B) *G. hirsutum* and C) *G. bickii*.

**Figure S3.**
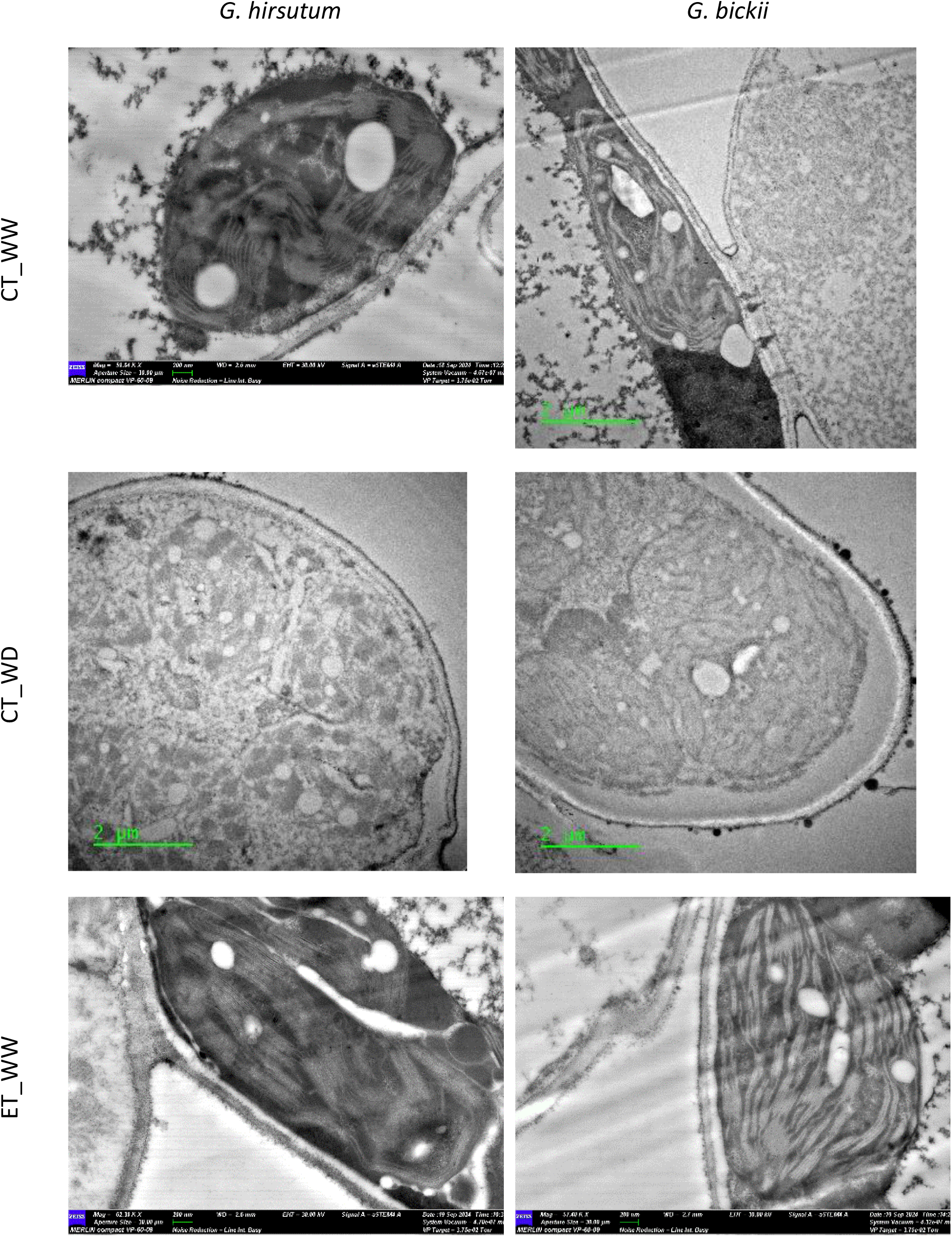
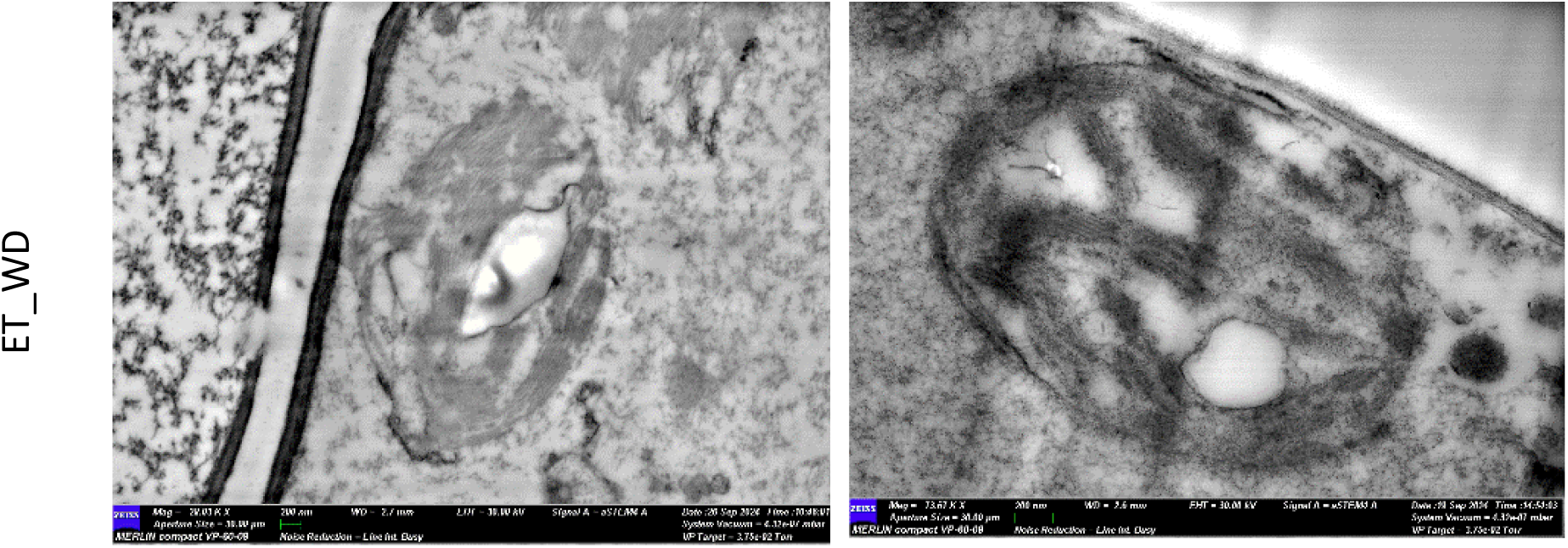
Transmission and scanning transmission electron micrographs of mesophyll cells from *Gossypium hirsutum* and *Gossypium bickii* under temperature and water treatments. Rows represent treatments: control temperature well-watered (CT_WW), control temperature water-deficit (CT_WD), elevated temperature well-watered (ET_WW), and elevated temperature water-deficit (ET_WD). Columns represent species (*G. hirsutum*, left; *G. bickii*, right). Micrographs were obtained using transmission electron microscopy (TEM) and scanning transmission electron microscopy (STEM) as described in Methods. Scale bars = 2µm for TEM and 200nm for STEM.

**Figure S4.**
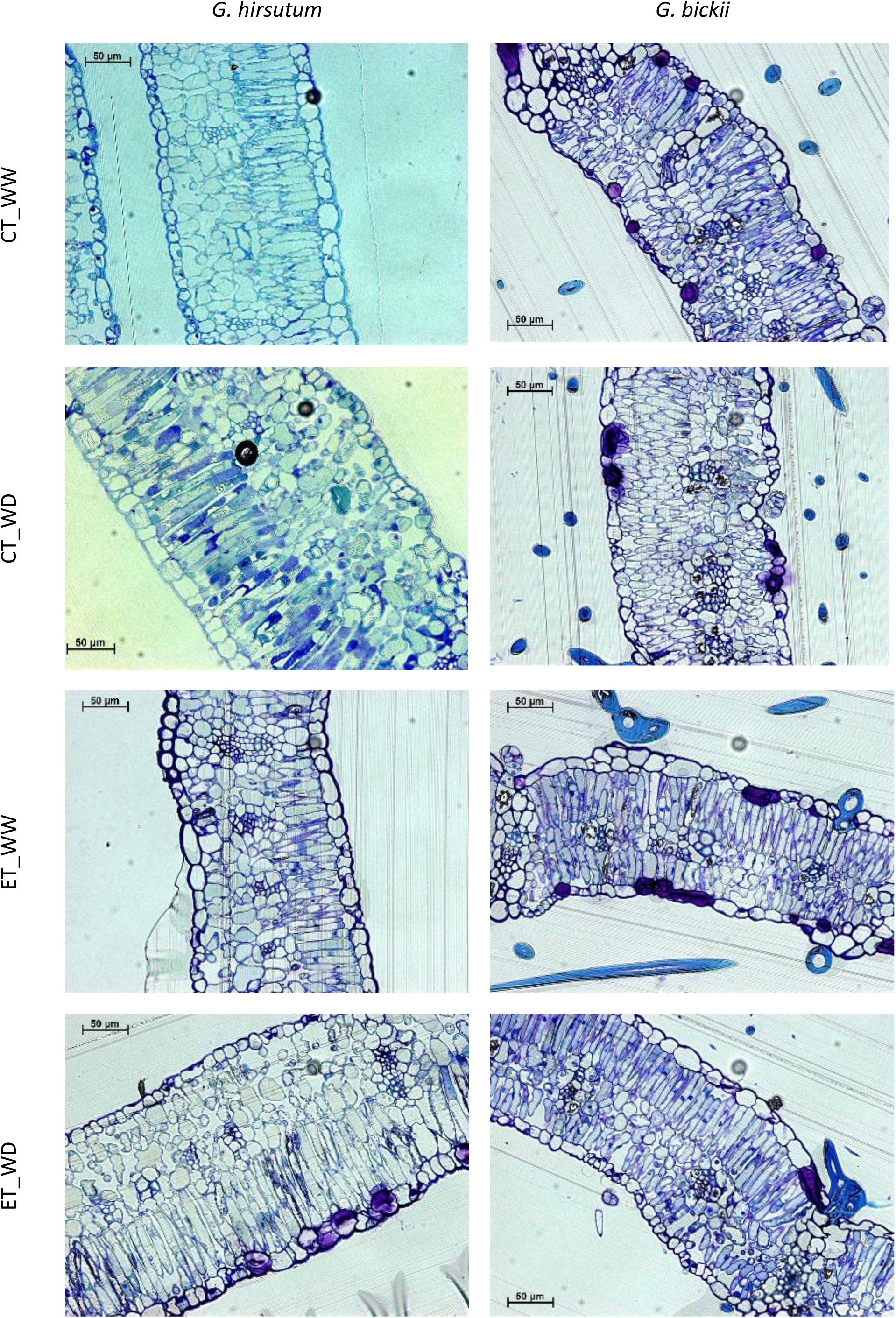
Optical micrographs of transverse leaf sections from *Gossypium hirsutum* and *Gossypium bickii* under temperature and water treatments. Rows represent treatments: control temperature well-watered (CT_WW), control temperature water-deficit (CT_WD), elevated temperature well-watered (ET_WW), and elevated temperature water-deficit (ET_WD). Columns represent species (*G. hirsutum*, left; *G. bickii*, right). Sections were stained with toluidine blue to visualise leaf anatomy, including epidermal layers, mesophyll tissues, and intercellular airspaces. Images were used for quantification of leaf thickness, mesophyll thickness, fraction of intercellular airspace (*F*ias), and mesophyll surface area exposed to intercellular airspace (*S*mes). Scale bars = 50 µm.

